# Rapid ordering of barcoded transposon insertion libraries of anaerobic bacteria

**DOI:** 10.1101/780593

**Authors:** Anthony L. Shiver, Rebecca Culver, Adam M. Deutschbauer, Kerwyn Casey Huang

**Affiliations:** Department of Bioengineering, Stanford University, Stanford, CA 94305, USA; Department of Genetics, Stanford University, Stanford, CA 94305, USA; Environmental Genomics and Systems Biology Division, Lawrence Berkeley National Laboratory, Berkeley, CA 94720, USA; Department of Plant and Microbial Biology, University of California, Berkeley, CA 94720, USA; Department of Microbiology and Immunology, Stanford University School of Medicine, Stanford, CA 94305, USA; Chan-Zuckerberg Biohub, San Francisco, CA 94158

**Keywords:** anaerobic bacteria, human gut microbiota, ordered mutant library, transposon insertion, barcode, Bar-seq, Tn-seq, RB-TnSeq, high-throughput genetics, systems biology

## Abstract

Commensal bacteria from the human intestinal microbiota play important roles in health and disease. Research into the mechanisms by which these bacteria exert their effects is hampered by the complexity of the microbiota and by the strict growth requirements of the individual species. The assembly of ordered transposon insertion libraries, in which nearly all nonessential genes have been disrupted and the strains stored as independent monocultures, would be a transformative resource for research into many microbiota members. However, assembly of these libraries must be fast and inexpensive in order to empower investigation of the large number of species that typically compose gut communities. The methods used to generate ordered libraries must also be adapted to the anaerobic growth requirements of most intestinal bacteria. We have developed a protocol to assemble ordered libraries of transposon insertion mutants that is fast, cheap, and effective for even strict anaerobes. The protocol differs from currently available methods by making use of cell sorting to order the library and barcoded transposons to facilitate the localization of ordered mutations in the library. By tracking transposon insertions using barcode sequencing, our approach increases the accuracy and reduces the time and effort required to locate mutants in the library. Ordered libraries can be sorted and characterized over the course of two weeks using this approach. We expect this protocol will lower the barrier to generating comprehensive, ordered mutant libraries for many species in the human microbiota, allowing for new investigations into genotype-phenotype relationships within this important microbial ecosystem.

## INTRODUCTION

There is a growing appreciation of the role that commensal bacteria play in human health^1^. However, the complexity of the microbiota, the fastidious growth requirements of its members^2^, and a lack of genetic tools and resources to study them have complicated efforts to unravel the mechanistic links between commensal bacteria and host health and disease. Ordered mutant libraries in which all non-essential genes have been comprehensively disrupted and individual mutant strains stored in monoculture have been invaluable resources for connecting genotype to phenotype, but such libraries exist for only a limited number of bacterial species^3–11^. This type of resource would be transformative for investigating the mechanistic basis for the effects of the gut microbiota on health; however, the generation of ordered mutant libraries must first be made rapid, inexpensive, and amenable to the anaerobic growth requirements of gut microbiota members.

Transposon mutagenesis is a highly successful mutagenesis approach for non-model organisms. A small number of well-characterized transposon systems (e.g., *mariner*^12^ and Tn*5*^13^) are functional in diverse bacteria, including the dominant phyla of the human microbiota. However, transposon insertion libraries are typically generated and assayed as a random pool. Many aspects of a bacterium’s interaction with a host cannot be readily studied in the context of a mutant pool, including phenotypes that can be complemented *in trans* such as the production of metabolites^14^. Thus, while transposon mutagenesis is a fruitful first approach for genetic analysis of under-studied gut microbiota members, a major challenge remains in ordering the random pool of transposon mutants in order to analyze individual strains.

Recent technological advances have vastly lowered the cost and effort required to order random transposon insertion libraries^15,16^. Pooling strategies that combine cultures prior to sequencing and then use computational methods to identify the positions of individual strains can reduce the number of samples that need to be processed by two orders of magnitude^5,7,10,16,17^. The strategy behind various pooling protocols can be separated into two parts. The first is pool design: how individual mutants are combined into larger pools to reduce the number of samples that must be processed, while maintaining the ability to deconvolve the locations of each mutant in the library. The second is how each pool is sequenced to identify the mutations contained within. Current sequencing approaches for transposon insertion libraries use a flavor of transposon insertion sequencing (Tn-seq/INSeq/ST-PCR). Because these sequencing approaches map transposon insertions for every individual pool, they remain laborious, often requiring multiple processing steps for each individual pool^7,10,15,16^.

We have incorporated two further advances to facilitate sorting of obligate anaerobic bacteria and further lower the cost and effort of locating mutations within an ordered library. First, we use single cell sorting with a fluorescence-activated cell sorting (FACS) machine to order individual strains. Second, we make use of barcoded transposons and barcode sequencing to locate individual mutations within the ordered library. The first advance makes use of the fast speed at which single cells can be isolated using a cell sorter (>5,000 mutants per hour) to order strains in high throughput while keeping aerobic exposure time below 30 minutes. The second advance utilizes the reproducibility and simplicity of barcode sequencing to reduce the effort required to locate mutations in the ordered library.

We recently developed the use of barcoded transposons and barcode sequencing (RB-TnSeq) as an alternative sequencing approach for high-throughput functional genomics^18^. With RB-TnSeq, the entire transposon insertion library is initially sequenced as one sample, allowing association of random DNA barcodes with transposon insertions across the entire pool. Subsequently, the abundance of transposon insertions can be followed with a single PCR step to amplify barcodes followed by next-generation sequencing (Bar-seq). This approach allowed us to meaure genome-wide mutant fitness data for 32 diverse bacteria across dozens of growth conditions^19^. The same principles that make Bar-seq useful for chemical genomics also make it an ideal sequencing approach for locating mutations within an ordered library.

While Bar-seq simplifies the process of locating mutants within an ordered library, it requires a diverse progenitor pool with a large number of unique barcodes. Therefore, the main limitation of our protocol is that it is restricted to bacterial strains with a relatively high transformation efficiency such that diverse pools of barcoded transposon insertions can be generated. If a high efficiency transformation protocol can be identified for a strain of interest, the use of Bar-seq promises to lower the barrier to generating an ordered library from a random pool of barcoded transposon insertions.

We have incorporated flow sorting and barcode sequencing into a protocol (Figure 1) that allows us to rapidly generate an ordered collection of >10,000 barcoded transposon insertion mutants in strictly anaerobic bacteria such as those found in the human gut microbiota. To develop this protocol, we first demonstrated that strict anaerobes can survive the temporary exposure to oxygen that occurs during single-cell sorting with a FACS machine, particularly in the presence of specific antioxidants. We capitalized on this finding to develop a protocol built around sorting single cells from a pooled mutant library. We optimized this protocol to reduce the time, cost, and levels of cross-contamination. Finally, we used this protocol to order a barcoded transposon insertion library in the model human gut commensal *Bacteroides thetaiotaomicron* VPI-5482. Analysis of individual mutants from this ordered library allowed us to further characterize chemical sensitivities that were detected from a pooled chemical-genomics screen of *B. thetaiotaomicron* VPI-5482^20^.

**Figure 1:**
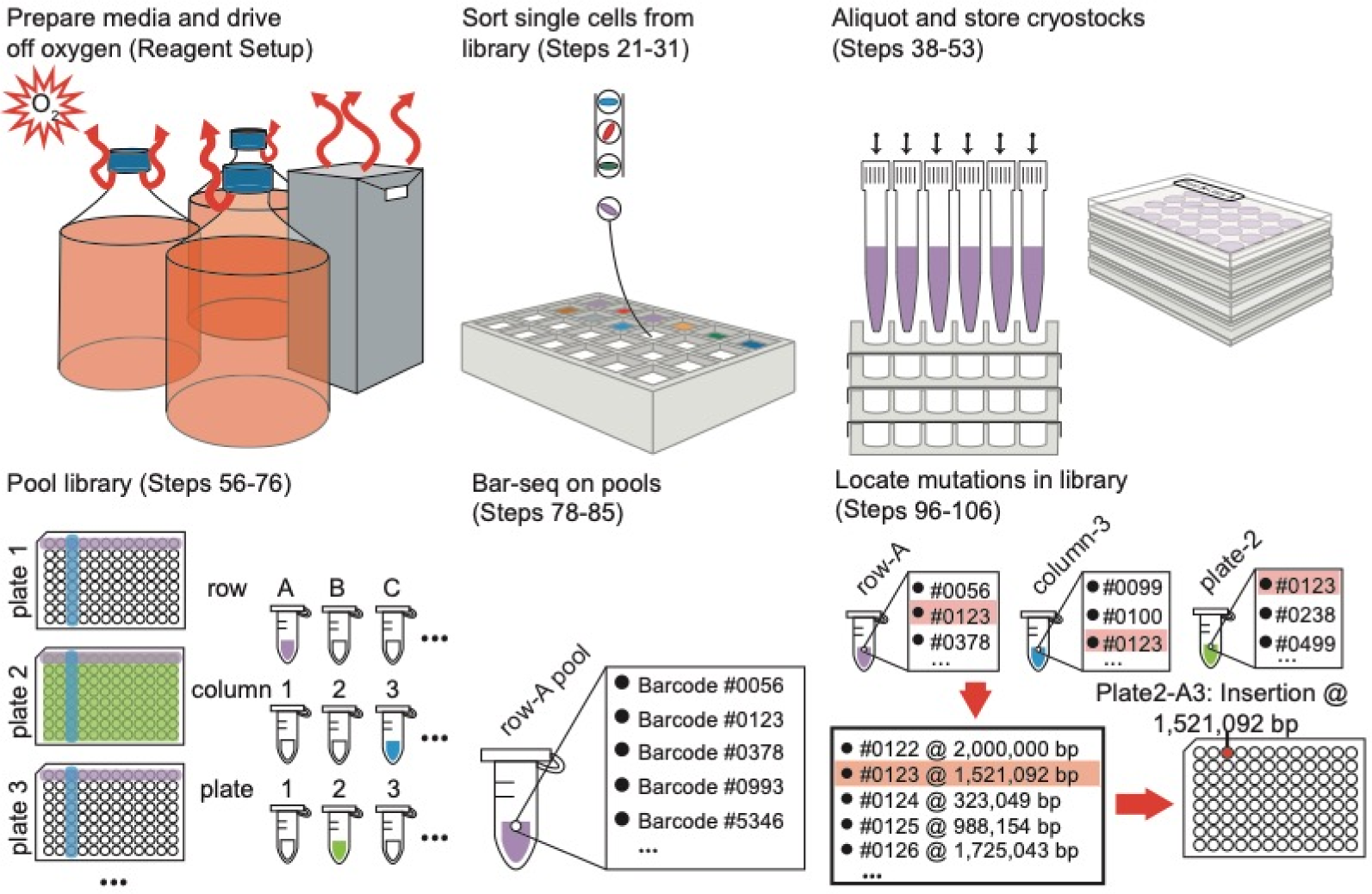
Schematic of the entire process of sorting the transposon-insertion library and mapping the barcodes. The protocol includes steps to order a random transposon library and steps to locate individual mutant strains in the library. (Reagent Setup) The media and consumables for a high-throughput library are made and allowed to sit in an anaerobic chamber long enough to drive off oxygen. (Steps 21-31) Single cells from a transposon mutant pool are sorted into individual wells of 96-deepwell plates and allowed to grow to saturation. (Steps 38-53) The cultures derived from sorted single cells are mixed with glycerol to make a cryostock, aliquoted into multiple copies, and stored at −80 °C. (Steps 56-76) Left-over cultures are pooled into subsets for sequencing. (Steps 78-85) Barcodes from the pools are PCR-amplified and sequenced. (Steps 96-106) Information on the inclusion of barcodes in each pool is used to locate barcodes within the library. Association of each barcode with a transposon insertion locates mutant strains within the library.

Given a diverse transposon insertion pool, both the sorting portion of the protocol and the strain identification portion can be accomplished in two weeks. As transformation methods are developed for more bacterial residents of the human microbiota, we expect that this protocol will enable the rapid and cost-effective investigation of microbe-host interactions, metabolite production, and other genotype-phenotype relationships in an ever-expanding set of bacteria.

## EXPERIMENTAL DESIGN

### Survival of anaerobic bacteria during sorting

To determine whether we could use cell sorting to generate ordered libraries of anaerobic bacteria, we first sought to quantify the ability of bacteria from the gut microbiota, particularly strict anaerobes, to survive the process of cell sorting. We chose two Bacteroidetes (*Bacteroides finegoldii* and *Parabacteroides johnsonii*), two Firmicutes (*Lactobacillus reuteri* and *Clostridium symbiosum*), and two Actinobacteria (*Bifidobacterium breve* and *Bifidobacterium longum*) to test survival of the sorting process; both *Bifidobacterium* species, *C. symbiosum, P. johnsonii*, and *B. finegoldii* are all obligate anaerobes while *L. reuteri* is a facultative anaerobe^21^. As a control, the facultative anaerobe *Escherichia coli* BW25113 was also sorted. Strains were grown and sorted into commercial media formulations of MRS (*B. breve* and *B. longum*), RCM supplemented with hemin, menadione, tryptophan, and arginine (*C. symbiosum*) and GAM supplemented with hemin, menadione, tryptophan, and arginine (*P. johnsonii*, *B. finegoldii*, *L. reuteri*, and *E. coli*).

All plates and cultures spent 1-1.5 h in an aerobic atmosphere while single cells were sorted into five 96-microwell plates for each strain. We counted the number of blank wells after outgrowth in each 96-well plate as a proxy for the number of single cells that did not survive the sorting procedure. We found that even the strict anaerobes survived the aerobic exposure long enough to complete the cell sorting protocol and regrow at high frequency when returned to anaerobic conditions. The fraction of blank wells after sorting varied across species, with *C. symbiosum* having the highest fraction of blank wells (**Figure 2a**); nonetheless, at least 89% of *C. symbiosum* cells grew after sorting.

**Figure 2:**
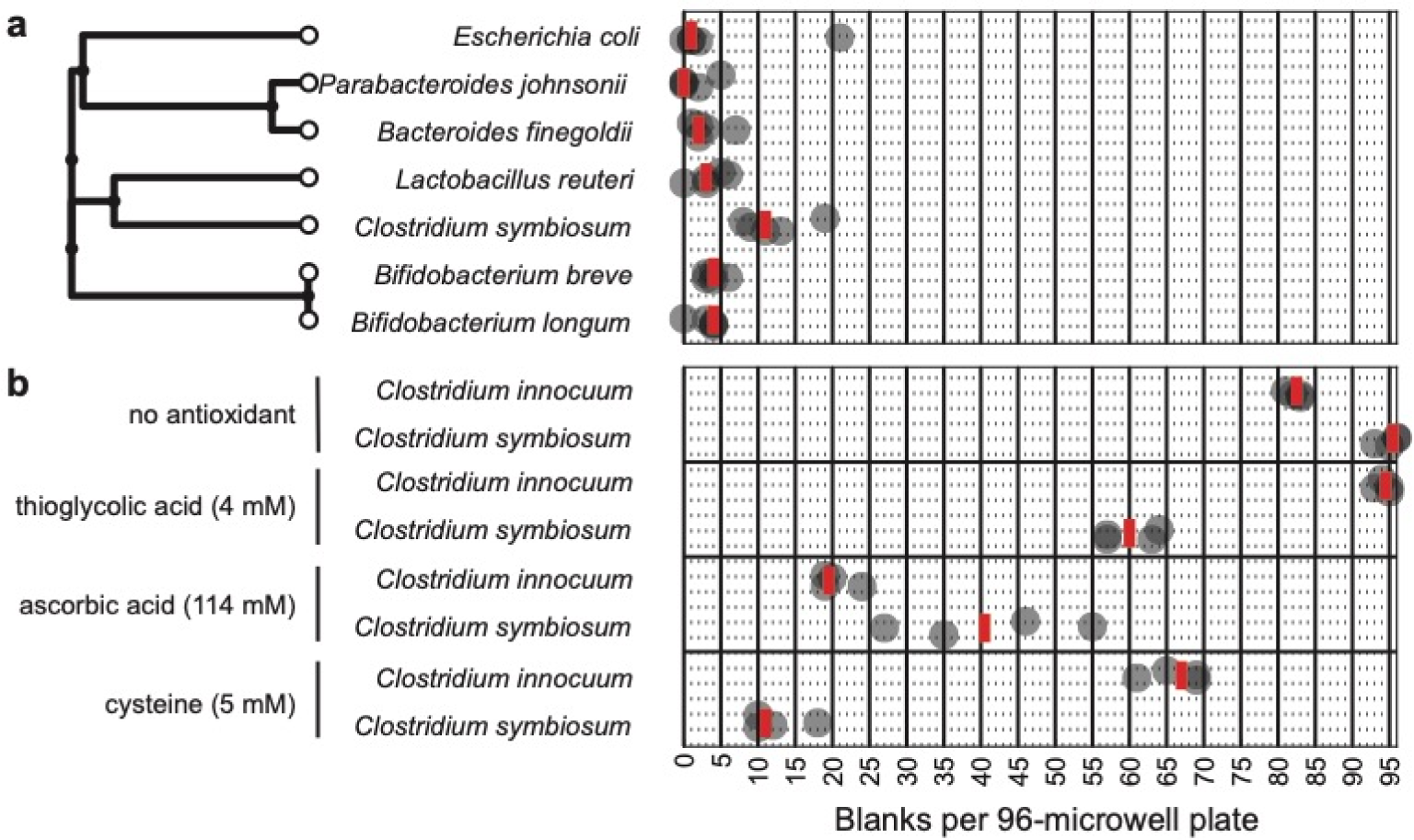
Strict anaerobes isolated from the human gut microbiota survive sorting in the presence of antioxidants. a) Six human gut isolates and a lab strain of *E. coli* survived sorting and grow in commercial media. Each strain was grown in a commercial formulation of media with added supplements. *Escherichia coli* BW25113, *Parabacteroides johnsonii* CL02T12C29, *Bacteroides finegoldii* CL09T03C10, and *Lactobacillus reuteri* CF48-3A were grown in Gifu Anaerobic Medium (GAM) supplemented with 1.5 µM hemin, 30 µM menadione, 6 mM arginine, and 1 mM tryptophan. *Clostridium symbiosum* WAL-14163 was grown in Reinforced Clostridial Medium (RCM) supplemented with 1.5 µM hemin, 30 µM menadione, 6 mM arginine, and 1 mM tryptophan. *Bifidobacterium breve* DSM20213 and *Bifidobacterium longum* NCC2705 were grown in de Mann, Rogosa, Sharp (MRS) medium. Each strain was sorted into five 96-microwell plates. Dark circles are the number of blank wells in one 96-microwell plate after outgrowth. Red lines are the median value for the 5 replicate 96-microwell plates. All strains except *E. coli* and *L. reuteri* are obligate anaerobes^21^. Phylogenetic distances were extracted from the TimeTree database^24^. b) Two highly oxygen-sensitive strains, *Clostridium symbiosum* WAL-14163 and *Clostridium innocuum* 6_1_30, were sorted and grown in media with an antioxidant. For both strains, we used an alternative formulation of RCM with no cysteine and no agar. We then added either no antioxidant, 4 mM thioglycolic acid, 114 mM ascorbic acid, or 5 mM cysteine. Dark circles are the number of blank wells in one 96-microwell plate after outgrowth. Red lines are the median value for the 4 replicate 96-microwell plates. It is possible to achieve high levels of viability for both of the strains.

Each of the commercial media formulations we used contained a significant concentration of cysteine, which we hypothesized was acting as an antioxidant to protect the cells during their transient exposure to oxygen. To test this hypothesis and explore the utility of alternative antioxidants, we sorted *C. symbiosum* and *Clostridium innocuum* in formulations of RCM in which cysteine had been either omitted or replaced with another antioxidant. We tested cysteine, thioglycolic acid (a component of GAM), and ascorbic acid for their ability to protect the two species. In the absence of any added antioxidants, the survival of both species was greatly reduced (Figure 2b), consistent with their sensitivity to oxygen. Both species were protected by the addition of antioxidants. Interestingly, *C. innocuum* preferred ascorbic acid over cysteine while *C. symbiosum* had the opposite preference (Figure 2b). Thioglycolate was the worst performing antioxidant of those tested for both species (Figure 2b). It remains to be determined if the optimal bacterium-antioxidant pair is idiosyncratic or if there is a general, as yet unidentified, protectant that can be applied across species. Nonetheless, it was straightforward to achieve high survival frequency for both of these strict anaerobes.

Multiple factors will be critical for successfully adopting the cell sorting protocol to a species of interest. First, the concentration of bacterium-sized inanimate particles must be low enough that sorting of these false signals does not appreciably reduce the fill rate of wells in the library. Filter sterilization of the growth medium using a 0.22-µm filter is effective for eliminating this source of error; however, particular species may produce these types of particles (e.g., outer membrane vesicles, extracellular polysaccharides) during the course of growth. Second, the majority of cells in the population must be viable. Third, single cells have to survive the time spent in an aerobic environment that is required to sort the library. Fourth, the majority of the cells in the population cannot exist as part of multi-cell aggregates. The first three issues will lower the fill rate of wells in the library, while the fourth problem will lower the fraction of wells in the library that contain a single strain.

Each of these potential issues is likely to depend on the growth conditions of the library. While extensive optimization of growth conditions for *B. thetaiotaomicron* was not necessary, there are a number of avenues for troubleshooting the sorting step for difficult bacteria. Changing the growth phase of the sorted cells (e.g. log phase versus early stationary phase) is likely to alter the fraction of cells that survive sorting. Testing different growth and recovery media may also be an important aspect of troubleshooting. Finally, while cysteine may need to be avoided as an antioxidant for known hydrogen sulfide producers (see below), there are other antioxidants that can be added as a supplement to the growth and recovery media.

### Elimination of H_2_S production

CAUTION Hydrogen sulfide is a toxic, caustic gas. Build-up of hydrogen sulfide inside the anaerobic chamber will degrade sensitive electronic equipment. Exposure to hydrogen sulfide gas should be minimized. Before working with large volumes of culture, determine the growth conditions that will prevent the production of hydrogen sulfide during growth.

Enzymes with cysteine desulfhydrase activity degrade cysteine and produce hydrogen sulfide, a toxic gas, as a byproduct. These enzymes are conserved across the eubacteria^22^, and *Bacteroides* sp. are known to produce hydrogen sulfide gas^21^. Given the large volumes of culture involved in growing an ordered a library, even moderate production of H_2_S has the potential to overcome standard H_2_S removal columns and pose a danger to personnel and equipment. Our original recipe for the BHIS growth medium of *Bacteroides thetaiotaomicron* included 8 mM cysteine as an antioxidant, and *B. thetaiotaomicron* cultures produced significant levels of hydrogen sulfide in pilot experiments.

To avoid the potential hazards of H_2_S production, we first confirmed that the level of hydrogen sulfide produced by *B. thetaiotaomicron* is proportional to the concentration of added cysteine. We held hydrogen sulfide test strips over 5 mL cultures of *B. thetaiotaomicron* with increasing amounts of cysteine. Reaction of hydrogen sulfide with lead acetate in the test strip led to the black lead sulfide product (**Figure 3a**). We then tested growth of *B. thetaiotaomicron* in BHIS as a function of added cysteine and found that cysteine could be left out of the BHIS recipe without adverse effects on anaerobic growth. The growth curve of *B. thetaiotaomicron* grown in BHIS without cysteine was similar to BHIS with cysteine added (**Figure 3b**). Calculation of the maximum growth rate and the maximum OD_600_ (optical density at 600 nm) also showed no significant differences between BHIS with or without added cysteine (**Figure 3c**). From these results and the known oxygen tolerance of *B. thetaiotaomicron*, we decided to proceed without cysteine for sorting the transposon insertion library.

**Figure 3:**
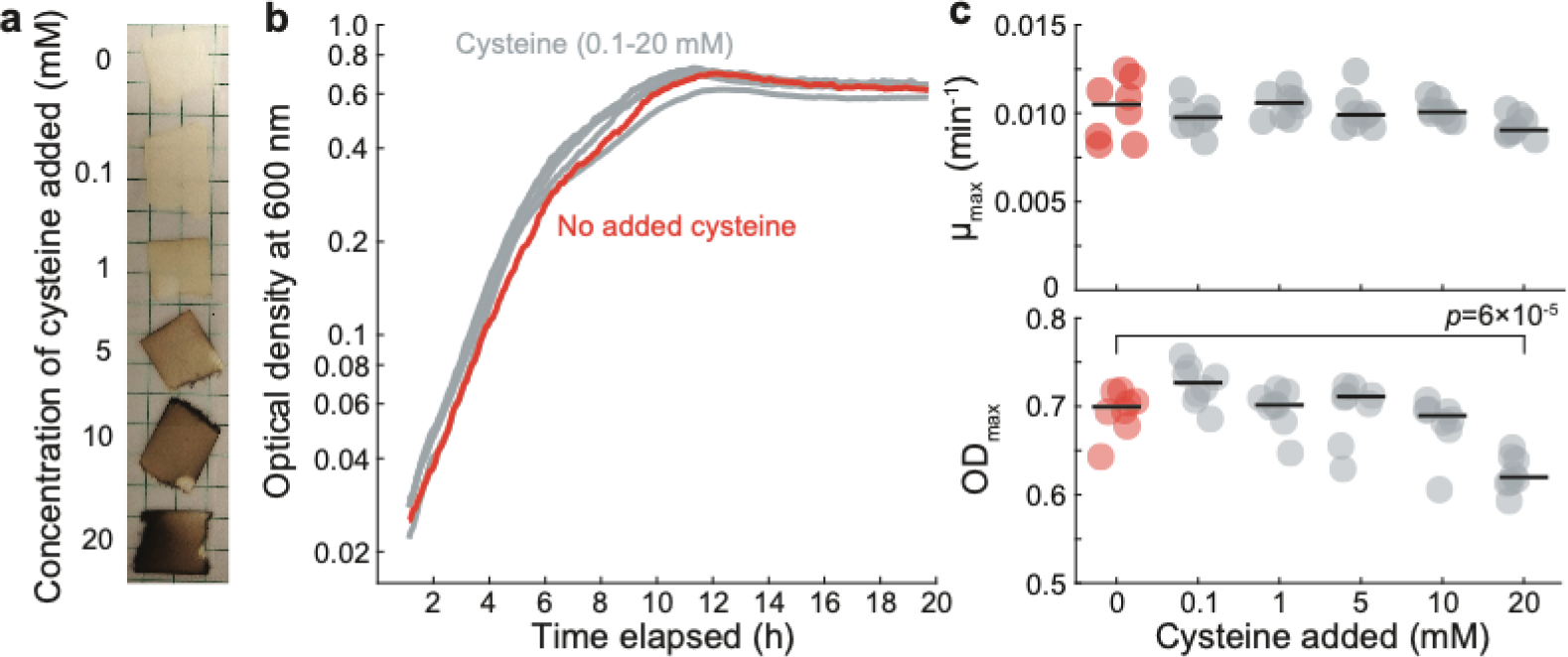
Elimination of H_2_S production may be necessary for large volumes of culture. a) *B. thetaiotaomicron* produces hydrogen sulfide proportional to the concentration of cysteine added to the medium. The original recipe for BHIS included supplementation with 8 mM cysteine. BHIS media with a range of cysteine concentrations were prepared and inoculated with *B. thetaiotaomicron*. After 24 h of growth, a hydrogen sulfide test strip was held over the culture for 5 s. Reaction of hydrogen sulfide with lead acetate in the strips leads to black discoloration. b) Anaerobic growth of *B. thetaiotaomicron* is not strongly impacted by the addition of cysteine. A 96-well microwell plate was filled with BHIS and increasing levels of cysteine (0-20 mM). The media were inoculated with *B. thetaiotaomicron* VPI-5482 and growth was monitored by reading optical density at 600 nm. The growth curves show the median value for (*n*=8) independent measurements. c) The maximum growth rate (μ_max_) (top) and maximum OD_600_ (OD_max_) (bottom) were extracted from individual growth curves (*n*=8). The individual data points (circles) and median (solid line) are plotted for each concentration of cysteine. The omission of cysteine (red circles) did not significantly alter the maximum growth rate or the maximum OD_600_. The only significant comparison between the no-cysteine condition and the cysteine-added conditions was with 20 mM cysteine for the OD_max_ parameter (multiple hypothesis corrected one-way ANOVA, *p*=6×10^−5^).

## MATERIALS

### REAGENTS

#### Media

- Bacto^TM^ Brain Heart Infusion (BHI) (Becton Dickinson, #237500)
- Sodium bicarbonate (Fisher Bioreagents, #BP328-500)
- Hemin (porcine) (Alfa Aesar, A11165)
- Erythromycin (Sigma, #E5389-5G)
- Glycerol (Fisher Bioreagents, #BP229-1)

#### Chemicals

- Sodium hydroxide (Fisher Bioreagents, #S318-500)
- Hydrochloric acid (Fisher Chemical, #A144-500)

#### Plasticware and Consumables

- Thermo Scientific^TM^ Nalgene^TM^ Rapid-Flow^TM^ sterile disposable bottle top filters with a PES membrane, 45-mm neck size (Thermo Scientific^TM^, #5974520) **CRITICAL** Polyether sulfone (PES) is superior to cellulose acetate (CA), surfactant-free cellulose acetate (SCFA), and polyvinylidene fluoride (PVDF) for filtering complex media like BHI without clogging.
- PlateOne 2-mL 96-deepwell polypropylene plates (USA Scientific, #1896-2000)
- Sterile 96-microwell V-bottom polystyrene plates (Grenier bio-one, #651161)
- Sterile microwell plate lids (Grenier Bio-One, #656101)
- Standard profile polypropylene reservoirs (Mettler-Toledo, # LR-R2-PB-5)
- Breathe-Easier gas permeable sealing membranes (Diversified Biotech, #BERM-2000)
- 5 mL polypropylene round bottom tubes (Falcon, #352063)
- Vinyl tape (3M, #471)
- Nunc Aluminum Seal Tape for 96-well plates (Thermo Scientific^TM^, #232698)
- Alumaseal 384 films (Excel Scientific, #F-384-100)
- 1-mL LTS filter tips, for use with BenchSmart (Mettler Toledo, #30296782)
- 200-µL LTS filter tips, for use with BenchSmart (Mettler Toledo, #17010646)
- 200-µL filter tips, for use with multichannel pipette (Rainin #30389239)
- 8.5” x 8.75” OD PAKVF4 MylarFoil pouches, opening on the ZipSeal end (Impak Corporation, #085MFSOZE0875)
- 500-cc oxygen absorbing packets (Impak Corporation, #SF500CS1500)
- Microtube tough-tags for laser printers, 0.91×0.32 in (Diversified Biotech, #9186-1000)
- Microtube tough-tags for laser printers, 1.28×0.5 in (Diversified Biotech, #TTLW-2016)

#### Glassware

- 2-L Pyrex bottles, 45-mm neck size (Pyrex, #1395-2L)
- 1-L Pyrex bottles, 45-mm neck size (Pyrex, #1395-1L)
- 250-mL Erlenmeyer flasks (Pyrex, #4980-250)

#### PCR and DNA isolation

- QIAamp 96 DNA QIAcube HT Kit (QIAgen #51331)
- 5x Q5 reaction buffer and high GC enhancer (New England Biolabs #B9027S)
- Deoxynucleotide (dNTP) Solution Mix (New England Biolabs #N0447L)
- Q5® High-Fidelity DNA Polymerase (New England Biolabs #M0491L)
- 96-well PCR plate (Eppendorf #0030133382)
- Zymo DNA clean and concentrator kit (Zymo Research #D4013)
- JumpStart DNA polymerase (Sigma Aldrich # D9307-50U)
- Agencourt AMPure XP beads (Beckman Coulter #A63880)
- Indexed primers for Bar-seq and RB-TnSeq (Integrated DNA Technologies; Table S1)
- NEBNext DNA Library Prep Master Mix Set for Illumina (New England Biolabs #E6040S)
- Covaris microTUBE AFA Fiber Pre-Slit Snap-Cap (Covaris #520045)

#### Oligos

- Order indexed primers (Supplementary Data 1) for amplifying the barcodes.
- Order oligos (Supplementary Data 2) for performing RB-TnSeq.

#### Strains

The sorting protocol requires a barcoded transposon insertion library that is compatible with Bar-seq. Briefly, a barcoded transposon insertion library is created using a traditional transposon mutagenesis approach (e.g. electroporation of *in vitro*-assembled transposomes or conjugation of the transposase and transposon on a non-replicating plasmid), with the addition on the transposon construct of a small sequence adjacent to one of the inverted repeats that acts as a barcode. In the approach from Ref. ^18^, the barcode is a stretch of 20 random nucleotides surrounded by two primer binding sites for amplification of the barcode. During the creation of a transposon insertion library, an RB-TnSeq experiment as described in steps 86-95 is performed to determine the size and quality of the library. The lookup table of insertion locations and the associated DNA barcodes generated by this RB-TnSeq reaction can be used to locate mutants within the ordered library using Bar-seq, or another RB-TnSeq reaction can be run using samples from the ordered library alone (as discussed below).

### EQUIPMENT

- Benchsmart 96 semi-automated pipetting system (installed within an anaerobic chamber), 1000 µL (Mettler Toledo, #BST-96-1000)
- Stylus Pen (Liberrway, #B01IHBVGOM)
- BenchSmart 96 pipette head, 200 µL (Mettler Toledo, #30296709)
- Forced air incubator for vinyl chambers (Coy Laboratory Products, #2000)
- Teflon-coated 6” hot jaw sealer, portable, constant heat (Impak Corporation, #IPKHS-606T)
- BD GasPak EZ large incubation container (Becton Dickinson, #260672)
- BD Influx Cell Sorter (Becton Dickinson, #646500) or equivalent
- Genesys^TM^ 20 visible spectrophotometer (Thermo scientific, #4001-000) or equivalent
- QIAcube HT system (QIAgen, #9001793) or equivalent
- PCR machine (Eppendorf Mastercycler X50a # 6313000018) or equivalent
- Pipet-Lite multichannel pipette L12-200XLS+ (Rainin # 17013810)
- MiSeq system (Illumina # SY-410-1003) or equivalent.
- HiSeq 4000 system (Illumina #SY-401-4001) or equivalent
- Covaris S220 Focused Ultrasonicator (Covaris #500217) or equivalent

### SOFTWARE

The software used for random barcode transposon-site sequencing is available from https://bitbucket.org/berkeleylab/feba/src. The scripts for resolving mutant positions within a ordered library are available from https://bitbucket.org/kchuanglab/resolve_barcode_position/src. For installation and configuration, follow the instructions in the respective repository.

### REAGENT SETUP

#### Bar-seq indexed primer plate

Combine Barseq_P1 with each unique, indexed reverse primer (BarSeq_P2) at a final concentration of 4 µM for each primer and store in a 96-well plate format. Store the oligo mixes at −20 °C, where they will be stable for years.

#### RB-TnSeq Y-adapter

Mix the oligos MOD2_TruSeq and Mod2_TS_Univ in a 50 µL volume in a PCR tube at a concentration of 10 µM for each oligo. Use a PCR machine to anneal the oligos using the “Anneal Y-adapter” program. Store the annealed oligos at −20 °C, where they will be stable for years.

#### Bar-seq PCR master mix (32.5 µL mix for a 50 µL PCR reaction)

Make the master mix fresh and store on ice until ready to use.

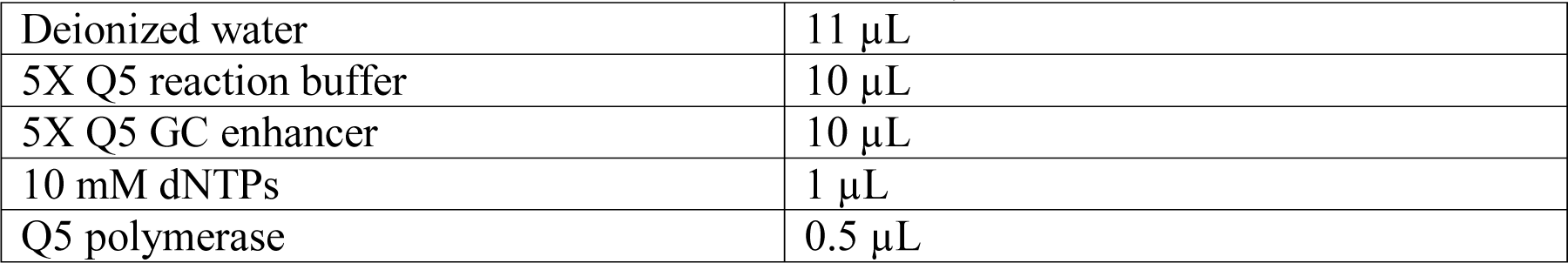

#### RB-TnSeq PCR master mix (50 µL mix for a 100 µL PCR reaction)

Make the master mix fresh and store on ice until ready to use.

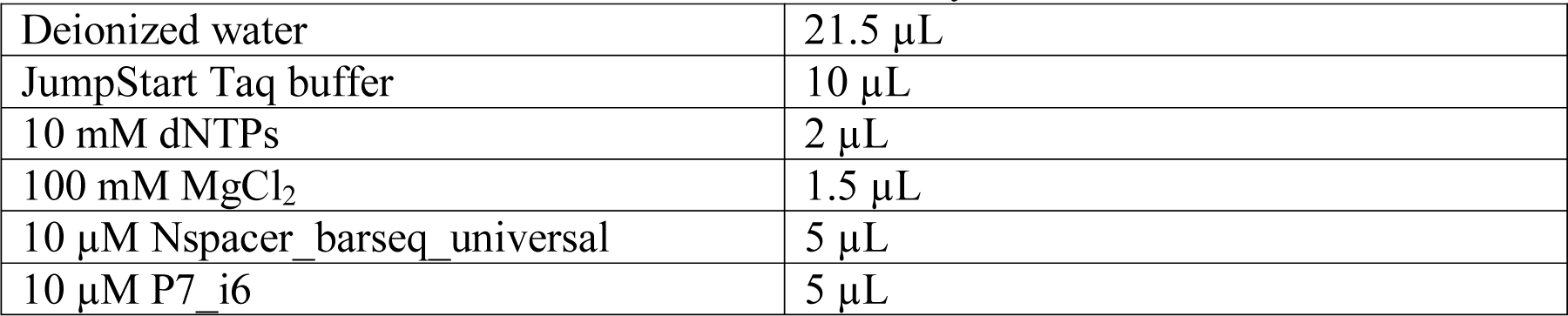
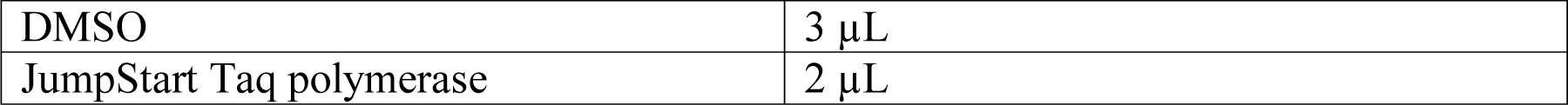

|  |  |
| --- | --- |
| Deionized water | 21.5 μL |
| JumpStart Taq buffer | 10 μL |
| 10 mM dNTPs | 2 μL |
| 100 mM MgCl<sub>2</sub> | 1.5 μL |
| 10 μM Nspacer_barseq_universal | 5 μL |
| 10 μM P7_i6 | 5 μL |
| DMSO | 3 μL |
| JumpStart Taq polymerase | 2 μL |

#### Media components

- Sodium bicarbonate (10% w/v, pH 7.4): To make 1 L, add 700 mL of deionized water to a 1-L beaker. Add 100 g of sodium bicarbonate and stir gently. Insert a calibrated pH meter and slowly add dilute HCl acid (0.1 M) to lower the pH to 7.4. When the target pH has been reached, add deionized water to a final volume of 1 L. Filter sterilize the solution with a 0.22-µm PES filter. The solution is stable for at least one month at room temperature with a tightly closed lid.

CRITICAL STEP Avoid heat, vigorous stirring, and concentrated acid solutions to prevent driving CO_2_ out of solution.

- Hemin (0.5 mg/mL in 0.01 M NaOH): To make 500 mL, add 250 mg of hemin to 5 mL of 1 M NaOH. Once the hemin has dissolved, bring the final volume to 500 mL with deionized water. Cover with foil and store at 4 °C until use. The solution is stable at 4 °C for several months.
- Antibiotic stock (e.g., 10 mg/mL erythromycin): Make an antibiotic stock at the desired concentration. For the *B. thetaiotaomicron* library, we dissolved 200 mg of erythromycin in 20 mL of absolute ethanol for a final concentration of 10 mg/mL, allowing us to make 20 L of BHIS supplemented with 10 µg/mL erythromycin. The stability of antibiotic stocks depends on the antibiotic and solvent. We stored the erythromycin stock at −20 °C and used the stock within a week.

#### Sterilization and pre-reduction of media and plasticware

The volume of media and number of plates have been chosen for a 40-plate library; adapt accordingly for larger libraries.

Make the sort recovery medium in 5 L batches: for BHIS, add 4 L of deionized water to a 5-L beaker. Add 185 g of commercial BHI mix and stir until the solution clears.

Microwave 500 mL of deionized water to bring to a boil and add to the solution, continue stirring. Add 50 mL of the hemin solution and 100 mL of the 10% sodium bicarbonate solution. Bring the final volume to 5-L using a graduated cylinder. Filter sterilize into 2-L flasks using a 0.22-µm PES filter. Repeat this process until all 15 L of medium have been made.

CRITICAL STEP It is critical to filter sterilize all media in which cells are grown prior to sorting in order to reduce the frequency of empty wells that result from sorting particulate matter. At a minimum, the 1 L of medium used to grow the library before sorting needs to be filter sterilized. We also filter sterilize the recovery medium to avoid wait time and batch effects due to the effects of autoclaving on the medium.

To prepare 3 L of 50% (w/v) glycerol, transfer 1.5 kg of glycerol to a 5-L beaker. Add water to 3 L final volume and transfer to 3 1-L bottles. Autoclave to sterilize.

Prepare and sterilize plastic and glassware: wrap 45 2-mL 96-deepwell plates in foil in stacks of 5. Autoclave to sterilize. Wrap 10 reservoirs in foil individually. Autoclave to sterilize. Cover the mouth of 4 250-mL Erlenmeyer flasks with foil. Autoclave to sterilize.

Move media and plasticware into the anaerobic chamber. Store the bottles of media and glycerol with bottle caps unscrewed to allow gas exchange. This includes:

- 7 2-L bottles of sort recovery medium
- 1 1-L bottle of sort recovery medium
- 2 sterilized reservoirs
- 45 sterilized 2 mL 96-deepwell plates
- 10 1-mL LTS filter tips
- 4 250-mL Erlenmeyer flasks
- 1 opened bag of 5 mL polypropylene round bottom tubes

Finally, let the media sit for 48-72 h in the anaerobic chamber.

CRITICAL STEP It is good practice to allow all materials that need to be anaerobic to sit in the anaerobic chamber for more than 48 h. If changes are made in the protocol to adapt for specific needs (e.g., duration of growth in step 32), then corresponding changes must also be made to when materials for the next set are brought into the anaerobic chamber to remove oxygen.

### EQUIPMENT SETUP

#### BenchSmart Programs

*Medium transfer*

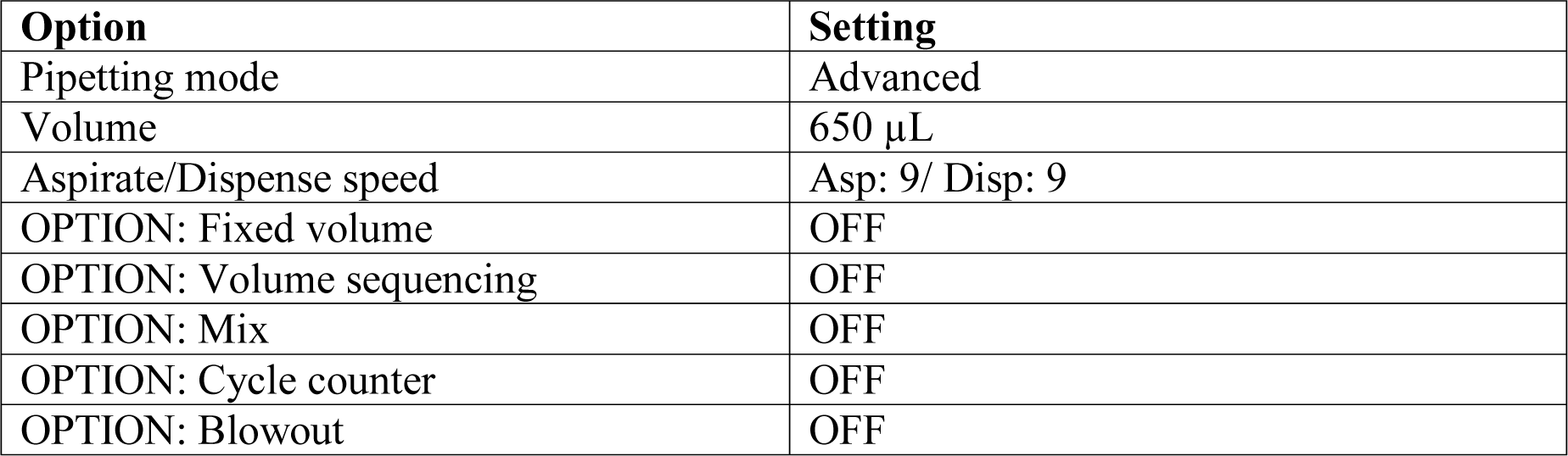

| Option | Setting |
| --- | --- |
| Pipetting mode | Advanced |
| Volume | 650 μL |
| Aspirate/Dispense speed | Asp: 9/ Disp: 9 |
| OPTION: Fixed volume | OFF |
| OPTION: Volume sequencing | OFF |
| OPTION: Mix | OFF |
| OPTION: Cycle counter | OFF |
| OPTION: Blowout | OFF |

*Dispense glycerol*

| Option | Setting |
| --- | --- |
| Pipetting mode | Advanced |
| Volume | 557 μL |
| Aspirate/Dispense speed | Asp: 5 / Disp: 5 / Mix: 5 |
| OPTION: Fixed volume | OFF |
| OPTION: Volume sequencing | OFF |
| OPTION: Mix | ON |
| OPTION: Cycle counter | OFF |
| OPTION: Blowout | OFF |
| MIX OPTION: Mix volume | 750 µL |
| MIX OPTION: Mix cycles | 9 |

*Repeat dispense*

| Option | Setting |
| --- | --- |
| Pipetting mode | MULTI-DISP |
| Aliquot volume | 100 µL |
| Number of aliquots | 1-10 |
| Aspirate/Dispense speed | Asp: 5 / Disp: 3 |
| OPTION: Fixed volume | OFF |
| OPTION: Volume sequencing | OFF |
| OPTION: Cycle counter | OFF |
| OPTION: Dispense blowout | OFF |
| OPTION: Auto pace | OFF |

*Mix Cultures*

| Option | Setting |
| --- | --- |
| Pipetting mode | Advanced |
| Volume | 200 µL |
| Aspirate/Dispense speed | Asp: 5 / Disp: 5 / Mix: 5 |
| OPTION: Fixed volume | OFF |
| OPTION: Volume sequencing | OFF |
| OPTION: Mix | ON |
| OPTION: Cycle counter | OFF |
| OPTION: Blowout | OFF |
| MIX OPTION: Mix volume | 200 µL |
| MIX OPTION: Mix cycles | 9 |

*Dispense pools*

| Option | Setting |
| --- | --- |
| Pipetting mode | MULTI-DISP |
| Aliquot volume | 15 µL |
| Number of aliquots | 2 |
| Aspirate/Dispense speed | Asp: 8 / Disp: 1 |
| OPTION: Fixed volume | OFF |
| OPTION: Volume sequencing | OFF |
| OPTION: Cycle counter | OFF |
| OPTION: Dispense blowout | OFF |
| OPTION: Auto pace | OFF |

*Combine pools*

| Option | Setting |
| --- | --- |
| Pipetting mode | Advanced |
| Volume | 100 µL |
| Aspirate/Dispense speed | Asp: 5 / Disp: 5 |
| OPTION: Fixed volume | OFF |
| OPTION: Volume sequencing | OFF |
| OPTION: Mix | OFF |
| OPTION: Cycle counter | OFF |
| OPTION: Blowout | OFF |

#### PCR programs

*Bar-seq*

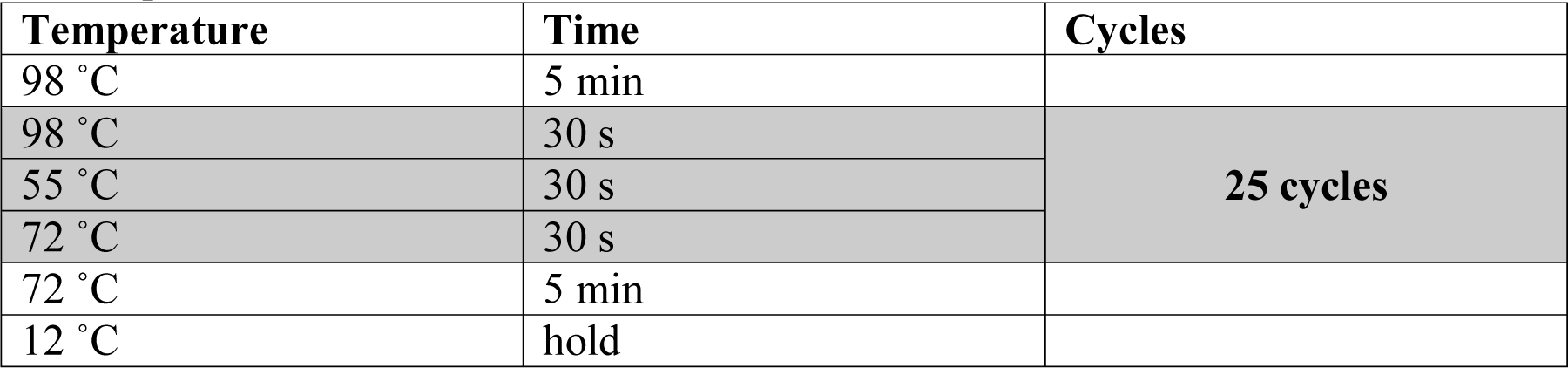

PCR programs *Bar-seq*
| Temperature | Time | Cycles |
| --- | --- | --- |
| 98 °C | 5 min |  |
| 98 °C | 30 s | 25 cycles |
| 55 °C | 30 s |  |
| 72 °C | 30 s |  |
| 72 °C | 5 min |  |
| 12 °C | hold |  |

*RB-TnSeq*

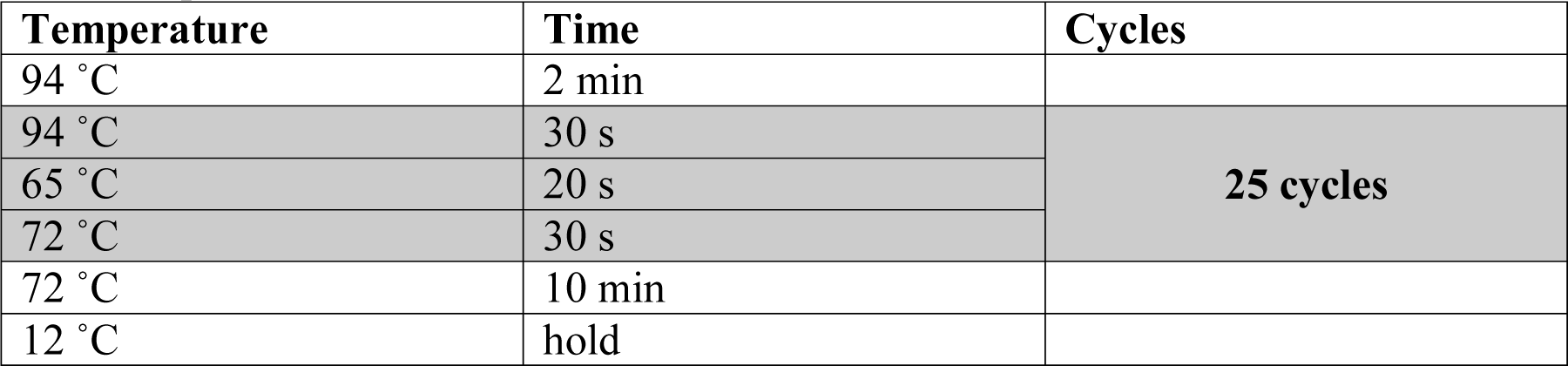

*Anneal Y-adapter*

| Temperature | Time | Cycles |
| --- | --- | --- |
| 95 °C | 2 min |  |
| 95 °C (-1 °C /cycle) | 38 s | 71 cycles |
| 12 °C | hold |  |

### PROCEDURE

#### Day 1: Prepare for the sort TIMING 0.5 d

1. (Optional) Remove shelving in the incubator to allow space for 2-mL 96-deepwell plates.
2. Add the selective antibiotic to the final concentration (e.g., for 10 µg/mL erythromycin: add 2 mL of 10 mg/mL erythromycin solution to every 2-L bottle of BHIS).
3. Load sterile 1-mL filter tips onto the BenchSmart. Use the tip box lid to cover a sterile empty reservoir.
4. Uncover the reservoir.
5. Add sort recovery medium to the reservoir up to 250 mL, if necessary.
6. Do two transfers of 650 µL (1.3 mL total) to a 2-mL 96-deepwell plate.
7. Cover the reservoir.
8. Seal the plate with an aluminum seal.
9. Repeat steps 4-8 until every 96-deepwell plate has been filled.
10. When the plates have been filled, unload the 1-mL tips. CRITICAL STEP Exercise caution when reusing the tips. Any contamination transferred to the tips through contact with a dirty surface will propagate throughout all subsequent plates. If you touch the tips unintentionally, eject them and load a fresh set before revisiting the medium in the reservoir.
11. Stack the 2-mL 96-deepwell plates inside the anaerobic 37 °C incubator.
12. Add the antibiotic to the final concentration to the 1-L bottle of sort recovery medium (e.g., add 1 mL of 10 mg/mL erythromycin to the 1-L bottle of BHIS).
13. Prewarm 4 250-mL Erlenmeyer flasks containing 100 mL sort recovery media with antibiotic.
14. One hour later, thaw a 2-mL aliquot of the mutant library and add the entire contents to one 250-mL flask (the other flasks will be used for maintaining the culture in log phase on the next day).
15. Grow the culture overnight at 37 °C.

#### Day 2: Sort the library TIMING 1 d

16. Determine the optical density of the overnight culture using a UV-Vis spectrophotometer. CRITICAL STEP Optical density measurements vary between plate reader machines, and every bacterial strain has an idiosyncratic growth curve in different media. Thus, it will be useful to run a growth curve using the same UV-Vis spectrometer used for this protocol to calibrate OD_600_ readings to the growth phase of your bacterium.
17. Dilute the overnight culture to an OD_600_=0.05 into one of the pre-warmed Erlenmeyer flasks of sort recovery media.
18. Periodically monitor the OD_600_, keeping the culture in log phase by diluting it into one of the pre-warmed Erlenmeyer flasks at a 1:4 dilution.
19. Pre-warm four 3-mL aliquots of sort recovery medium in FACS tubes.
20. Before sorting, measure the OD_600_ of the culture and dilute into the FACS tubes to cover a range of OD_600_ values (typically 0.01-0.05).
21. Tape the caps to the sides of the FACS tubes using vinyl tape to slow the entrance of oxygen.
22. Transfer the FACS tubes to a GasPak EZ container and bring them out of the anaerobic chamber along with the first set of 2-mL 96-deepwell plates.
23. Transport the material to the FACS machine, being careful not to agitate the plates.
24. Calibrate the cell sorter to ensure droplets are properly sorted into each well. Open the GasPak EZ container, vortex the first tube with the vinyl tape still on. Remove the tape and load the tube on the sample injection port. CRITICAL STEP After unsealing the tube, agitate the sample as little as possible. Without agitation the oxidation front will remain near the top of culture for longer, delaying the onset of oxidation for the cells entering the sample injection port at the bottom of the tube.
25. Calibrate the FACS machine to sort single cells: adjust the threshold and gain for forward scatter (FSC) and side scatter (SSC) so that the population of single cells has a clear distribution of signal. Define a restrictive two-dimensional gate on FSC and SSC that selects for the average signal from single cells and excludes larger signals. Use a secondary gate on pulse trigger width to further select for single cells. ?Troubleshooting CRITICAL STEP The gating strategy used to identify single cells is critical for successfully sorting the library and will depend heavily on the strain, growth medium, and physiological state of the cells. Test the exact conditions used to grow the library beforehand to determine the expected signal during the library sort. Unseal a 96-deepwell plate and cover with a sterile lid; check for liquid on seal. ?Troubleshooting
27. Sort single cells into the plate using the FACS machine.
28. Reseal the plate with a Breathe-Easier gas permeable membrane. Use a roller to ensure that a tight seal has formed, especially around the edges of the plate. CRITICAL STEP Ensuring a good seal is important to prevent evaporation from the outer rows and columns of the 96-deepwell plate during the recovery period. CRITICAL STEP It is important to avoid jostling or agitating the plates from this point onwards to minimize cross contamination between the wells. If a plate is jostled while applying the seal or moving to the incubator, check for wetting of the membrane and discard the plate if media has splashed onto the seal.
29. When the batch of plates has been sorted, return the plates to the anaerobic chamber and place them in the 37 °C incubator. CRITICAL STEP Cycle the airlock with a low-pressure program to avoid peeling the seal from the plates (e.g., 9 cycles of 5 in Hg vacuum pressure and fill with N_2_, followed by 9 cycles of 5 in Hg vacuum pressure and fill with the chamber atmosphere).
30. Take the next batch of plates out of the 37 °C incubator for sorting. CRITICAL STEP Bring the plates out of the anaerobic chamber in small batches. Optimize the size of each batch so that the plates spend as little time in the aerobic environment as possible.
31. Repeat steps 26-30 until cells have been sorted into all 96-deepwell plates (Figure 4).
32. Once all plates have been sorted, incubate them at 37 °C to allow recovery and growth of the sorted cells. ?Troubleshooting

**Figure 4:**
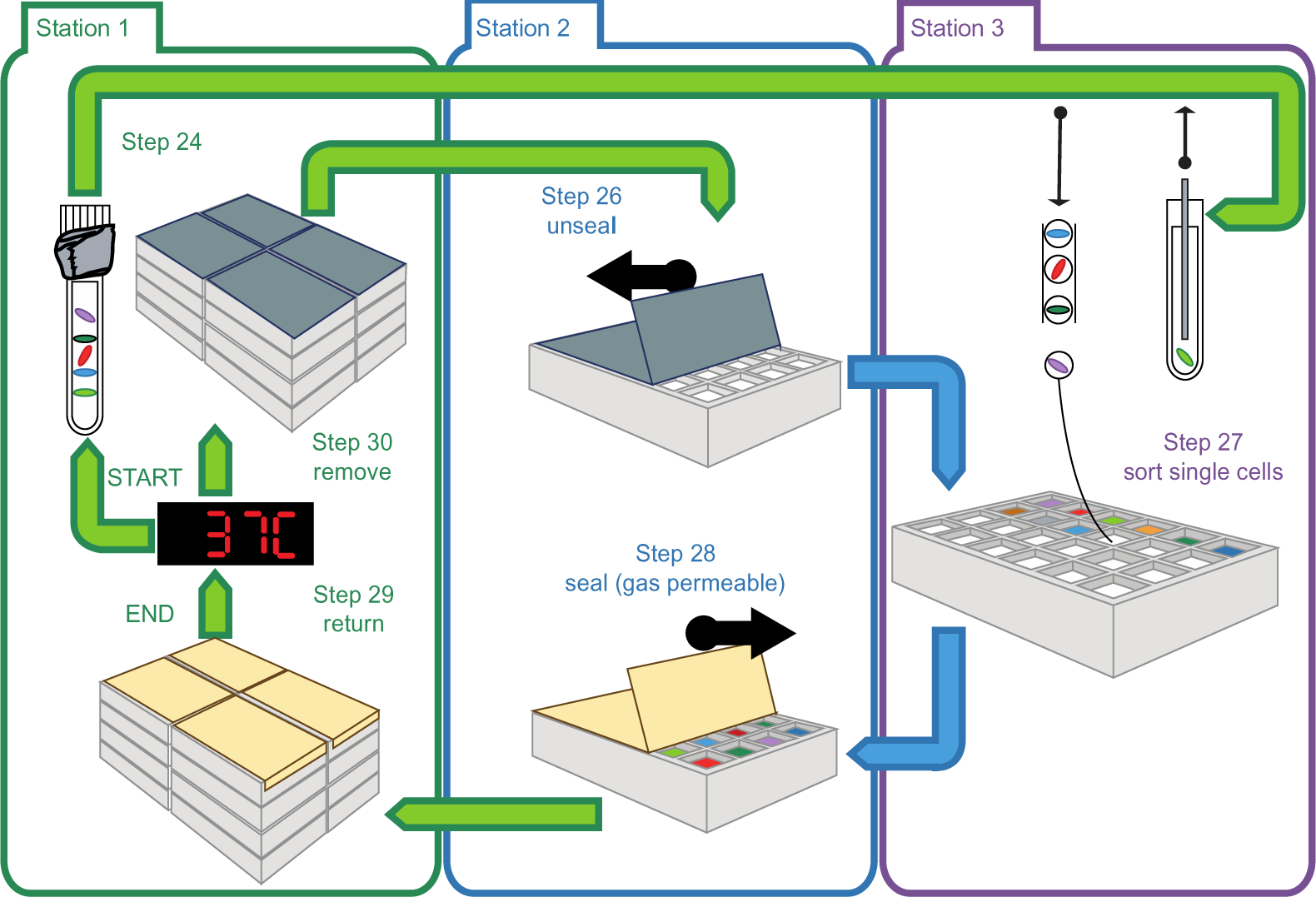
Sorting schematic. Three people work together to rapidly sort single cells. Station 1 is centered at the anaerobic chamber. The person working at station 1 cycles between the anaerobic chamber and the cell sorter, bringing sets of fresh (unsorted) 96-deepwell plates to the cell sorter and taking back the set of sorted plates that have accumulated. Station 2 is next to the cell sorter. This person works to unseal fresh plates, pass them to station 3, take sorted plates, and reseal them with a gas permeable membrane. Station 3 is the cell sorter. This person works to set up the equipment for the sort and run the single-cell sorting protocol. The arrows indicate movement of consumables between stations. The arrows are color coded by the person doing the work.

#### Day 3: Label and pre-reduce microwell plates TIMING 1 d

33. Unpack the 96-microwell plates.
34. Add a label to the front and to one side.
35. Add a sterile lid with a label, store in large sterile plastic bags.
36. Move the bags into the anaerobic chamber to drive oxygen off of the plastic. For 6 copies of a 40-plate library, process 240 96-microwell plates.
37. Move the appropriate number of MylarFoil bags into the anaerobic chamber, equal to the number of processed 96-microwell plates.

#### Day 4/5: Aliquot the library TIMING 1 d

38. Remove the Breathe-Easier membrane from a culture plate, cover with a sterile tip-box lid. CRITICAL STEP Take a moment to examine the growth pattern within the 96-deepwell plate. If the pattern is troubling (no growth, uniform growth in all wells) then discard the plate before it is assigned a number in the following step.
39. Add a label to the plate to assign a plate number. CRITICAL STEP This step will be the first point at which the culture plates are identified with a unique label. From this point onwards, it is important to check that the labels match when mixing, aliquoting, and storing the glycerol stock.
40. Transfer the culture plate to the BenchSmart, match with the correspondingly labeled 96-microwell plates.
41. Load 1-mL filter tips.
42. Switch the BenchSmart program to “Dispense glycerol”.
43. Transfer 557 µL of 50% glycerol to the cultures and mix. CRITICAL STEP Track the height of the culture when resuspending to keep the tips of the pipette just below the surface. Submerging the pipette tips when the wells are almost full will push the cultures out of the wells, while allowing the tips to dispense above the culture surface will cause splattering.
44. Switch the BenchSmart program to “Repeat dispense”.
45. Dispense 100 µL to each 96-microwell plate.
46. Unload tips.
47. Seal the 96-microwell plate glycerol stocks with an AlumaSeal. Use a roller to ensure that a good seal is formed.
48. Place the sealed microwell plates in MylarFoil heat seal bags along with a 500-cc oxygen absorber, zip-seal the bags.
49. Seal the original culture with an aluminum seal.
50. Once enough bagged plates have accumulated, move them outside the anaerobic chamber and use the heat sealer to seal the bags.
51. Move the bagged plates to the −80°C freezer for long-term storage.
52. When finished, or as needed to free up space, move the sealed original cultures out of the anaerobic chamber.
53. Repeat steps 38-52 until all 96-deepwell plates have been copied as cryostocks (Figure 5). CRITICAL STEP Aliquoting the library will require a team of people to work efficiently. We have designed this protocol around an anaerobic chamber with two sets of gloves to include a team of three researchers. One person will unseal, label and reseal the plates. One person will operate the BenchSmart to mix and aliquot the glycerol stocks. One person will work outside the anaerobic chamber to heat-seal and store the glycerol stocks at −80 °C (Figure 5).
54. At the end, transfer 200 µL of each of the original culture plates to a flat bottom 96-microwell plate and measure the OD_600_ using a plate reader. This measurement of the final OD_600_ of every well in the library will allow for an estimate of the fraction of wells in which growth occurred. Store the remainder of the culture at 4 °C or −20 °C. PAUSE POINT The remainder of the cultures in 96-deepwell plates will be used in subsequent steps to pool the library and sequence. The 96-deepwell plates can be stored overnight at 4°C or for a few days at −20°C.
55. Analyze the final OD_600_ values of the ordered library. Use a cutoff to determine the fraction of wells in which growth was detected. Examine the distribution of growth/no growth and final OD_600_ values across plates. Plates with abnormal growth can be triaged. ?Troubleshooting

**Figure 5:**
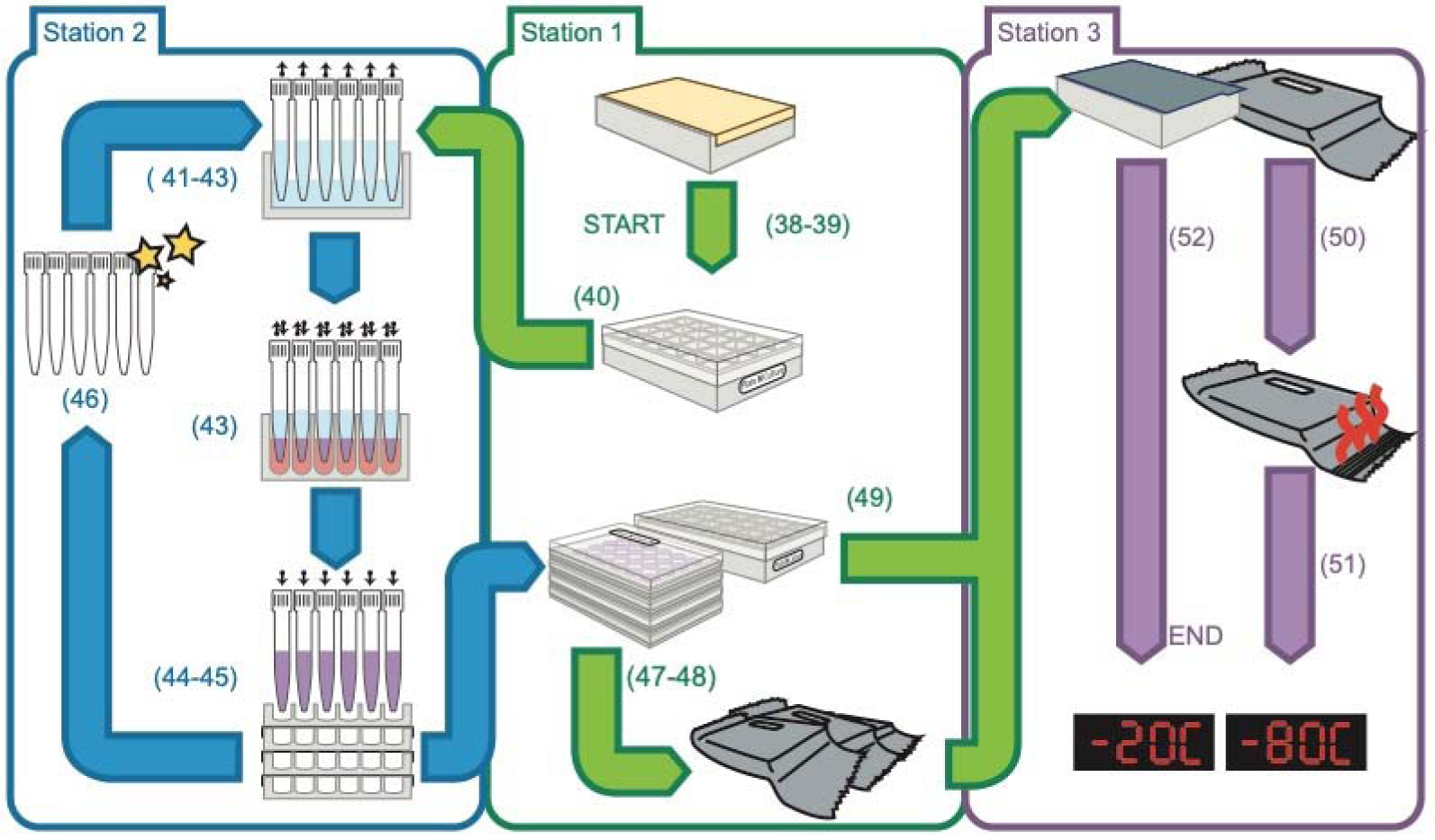
Cryostocking schematic. Three people work together to rapidly make and aliquot cryostocks of the library. Station 1 is at the incubator inside the anaerobic chamber. This person works to unseal the 96-deepwell plates, label them, and pass the plates to Station 2. Station 2 is at the BenchSmart inside the anaerobic chamber. This person works to mix the cultures with glycerol and aliquot the cryostocks into microwell plates. After aliquoting a set of cryostocks, this person passes them back to Station 1. Station 1 then seals the cryostocks with an aluminum seal, places them in a MylarFoil heat seal bag, and passes the cryostocks through the airlock to Station 3. Station 3 works next to the anaerobic chamber airlock. This person takes MylarFoil bags, heat-seals them, and stores the cryostocks at −80 °C.

#### Pool the library and extract genomic DNA TIMING 1d

56. Thaw the original 96-deepwell plates from step 54.
57. Centrifuge each 96-deepwell plate for 5 min at 4,500 x *g* at room temperature. CRITICAL STEP Prevention of cross-contamination during pooling is essential to accurately resolve the position of barcodes within the library. Centrifugation pulls down any culture that may be stuck to the seal and prevents droplets from spraying between wells when the seal is pulled off.
58. Allow the 96-deepwell plates to thaw at room temperature.
59. Open two sterile 2-mL 96-deepwell plates and cover each one with a sterile tip-box lid. The first 96-deepwell plate will store a mixture of each well from all plates (all-sample plate). The second 96-deepwell plate will store the final pools (final-pool plate). CRITICAL STEP The following steps are designed to work with a 40-plate library so that individual pools have a volume of less than 1.5 mL. To adapt for a larger library, lower the volumes accordingly. CRITICAL STEP The following steps are designed to work with a pool design and library size that need less than 96 pools. If more than 96-pools are required, more than one final-pool plate will be necessary. CRITICAL STEP Mislabeling or cross-contaminating wells in the final-pool plate will lead to significant errors in locating barcodes. Work carefully to ensure that no pipetting errors are made when filling the plate.
60. Load sterile 200-µL filter tips onto the BenchSmart.
61. Set the BenchSmart-96 to the “Mix cultures” program.
62. Aspirate from the centrifuged 96-deepwell plate and dispense into the same wells to mix the cultures. CRITICAL STEP If the cultures are prone to settling, it is important to dispense aliquots into the pools quickly to avoid pipetting errors.
63. Set the BenchSmart-96 to the “Dispense pools” program.
64. Remove the seal from the centrifuged 96-deepwell plate and cover with a sterile tip box lid.
65. Open a sterile, clean, single-well reservoir.
66. Multi-dispense 15 µL into the single-well reservoir, creating spots on the bottom of the reservoir. CRITICAL STEP Keep the tips elevated from the bottom of the reservoir to prevent backup of the dispensing liquid, which can lead to spattering of the liquid and cross contamination between tips.
67. Multi-dispense 15 µL into the all-sample plate.
68. Collect the culture from the single-well reservoir and store in a well of the final-pool plate.
69. Repeat steps 64-68 for every plate in the library.
70. Pool the library according to rows and columns using the all-sample plate.
71. For every row in the all-sample plate, use a 12-channel 200-µL pipette with 12 filter tips loaded. Transfer 120 µL per well to a single-well reservoir. Collect the culture and store in a well of the final-pool plate.
72. For every column in the all-sample plate, use a 12-channel 200-µL pipette with 8 filter tips loaded. Transfer 180 µL per well to a single-well reservoir. Collect the culture and store in a well of the final-pool plate. OPTIONAL: Steps 73-76 are included to generate DNA for running RB-TnSeq on the ordered library.
73. Load 200-µL filter tips onto the BenchSmart.
74. Set the BenchSmart-96 to the “Combine pools” program.
75. Use the BenchSmart-96 to transfer 100 µL from the all-sample pool plate to a single-well reservoir. Transfer the combined sample into a 15-mL conical tube then distribute the sample in 1.5 mL aliquots across multiple wells of the final-pool plate.
77. Extract genomic DNA from the final-pool plate following the manufacturer’s protocol for the QIAamp 96 DNA QIAcube HT kit.

#### Perform Bar-seq to identify the pool inclusion patterns of barcodes TIMING 1d

78. Make enough of the PCR master mix to amplify each pool, add 32.5 µL per reaction to the wells of a 96-well PCR plate.
79. Add 5 µL of the 4 μM indexed primer mixes to each well.
80. Add 12.5 µL of the extracted genomic DNA to each well.
81. Seal the PCR plate with a foil seal and run the Bar-seq PCR program.
82. Run 10 µL of each PCR product on a 1% (w/v) agarose gel to check for the correct band (∼200 bp).
83. Combine 5 µL of each Bar-seq product into a single pool. Store the remainder.
84. PCR purify a 50 µL aliquot of this total-library pool by following the manufacturer’s protocol for a Zymo DNA clean and concentrator kit. Elute in 20 µL of sterile deionized water.
85. Submit the purified PCR product for sequencing using a HiSeq 4000, 1×50 bp reads.

#### OPTIONAL: Perform RB-TnSeq to associate barcodes with insertion sites TIMING 1d

Performing RB-TnSeq on the ordered transposon library in addition to the complete insertion pool is not necessary but may be helpful in certain cases. If the same barcode is associated with more than one insertion site, then the true nature of this insertion is ambiguous: two independent strains, with one insertion per genome each, could serendipitously share the same barcode; alternatively, if the library was generated using conjugation, then repeated barcodes may arise from multiple transposon insertions in the same genome. If the initial insertion pool is complex and has a high fraction of ambiguous barcodes, then performing RB-TnSeq on the ordered library will likely reduce the fraction of ambiguous barcodes since it is improbable that two strains carrying the same barcode will both be sorted.

86. Combine genomic DNA aliquots from the whole-library pool of step 76 and dilute so that there is 1.5-3 µg DNA per aliquot.
87. Fragment the purified genomic DNA to an average size of 300 bp using a Covaris S2 Ultrasonicator using the manufacturer’s protocol.
88. Further size-select the fragmented DNA for an average size of 300 bp with a tighter distribution using AMPure XP beads using the manufacturer’s protocol.
89. At the adapter ligation step, ligate the custom TruSeq Y*-adapters (Reagent Setup) to the DNA fragments at a final concentration of 150 nM. In the clean-up step after adapter ligation, elute into 50 µL of elution buffer to use in the next PCR step.
90. Make enough of the PCR Master mix for a 100 µL reaction (See Reagent Setup).
91. Add 50 µL of adapter-ligated DNA.
92. Run the RB-TnSeq PCR program.
93. Run 5 µL of PCR product on a 1% (w/v) agarose gel to check for the correct band (200-300 bp).
94. Purify the PCR product with AMPure XP beads.
95. Submit the purified PCR product for sequencing using a MiSeq, 1×150 bp reads.

#### Locate the transposon insertion mutants in the library TIMING 1d

96. Create a new directory and assemble the necessary input files inside.
97. Download the appropriate genbank file with gene annotations.
98. Convert the genbank file into *fasta* and *genes* files for further analysis (Figure 6) using the command “*SetupOrg.pl -gbk genbankfile -out organism*”.

SetupOrg.pl generates a *fasta* file for the genome and a *genes* tab-delimited file that are used as inputs in subsequent analysis.
99. Move the RB-TnSeq fastq file(s) into a directory named *tnseq*.
100. Move the Bar-seq fastq files into a directory named *barseq*.
101. Create or move a model file into a directory named *model*.
  The model file is a model sequence for what should be expected before the transposon-genome junction. Run “*MapTnSeq.pl -h*” for further explanation of how the model file is constructed and see *feba/primers/model_\** in the FEBA repository for examples.
102. Map reads onto the genome (Figure 6) by running the command *MapTnSeq.pl - genome organism/genome.fna -model model/modelfile -first tnseq/fastq > tncount*
103. Generate the lookup table (Figure 6) by running the command *python3 path_to_repository*/*scripts/DesignSortedLibrary.py3 --input tncount --genes organism/genes.tab --Nmin 10 --output lookuptable*
104. Generate a tab-delimited key file (*barseqkey,* Figute 6) with the columns *path_to_fastQ file*, *index name*, and *pool name*.
  The FEBA codebase uses a restricted set of indexes to identify the barcodes in a Bar-seq read. The index name in the key file indicates the index that should be used for a demultiplexed fastq file using the FEBA set of primers. When using an alternative set of primers for Bar-seq, the scripts will have to be modified accordingly. This key file allows for the association of each sequencing file with the appropriate pool label. The pool names must be P# for plate number, C# for column, and R# for row, e.g. P1, C3, or R5.
105. Generate the pool inclusion table (Figure 6) by running the command *python3 path_to_repository*/*scripts/SortedPoolCounts.py3 –k barseqkey –o inclusiontable –q 10 –f 0.01* ?Troubleshooting CRITICAL STEP Unlike the other sequencing approaches, which map reads onto the genome, there is no internal metric that indicates whether a given barcode sequence is real or spurious. Prudently choosing the minimum number of reads to consider a barcode real is important to avoid introducing extra barcode sequences into the analysis pipeline. *SortedPoolCounts.py3* uses the *-f* parameter, the minimum fraction of reads relative to expectation, to control the number of barcode sequences included in the analysis.
106. Locate transposon insertion mutants in the ordered library (Figure 6) by running the command *matlab -nodesktop -nosplash -r \*
  *“addpath(path_to_repository/scripts/); \* resolve_barcode_position(inclusiontable,lookuptable,organism/genes.tab,positiontable); \ *exit;”*
107. Examine the *positiontable* file; this is the final map of strain locations within the library. ?Troubleshooting

**Figure 6:**
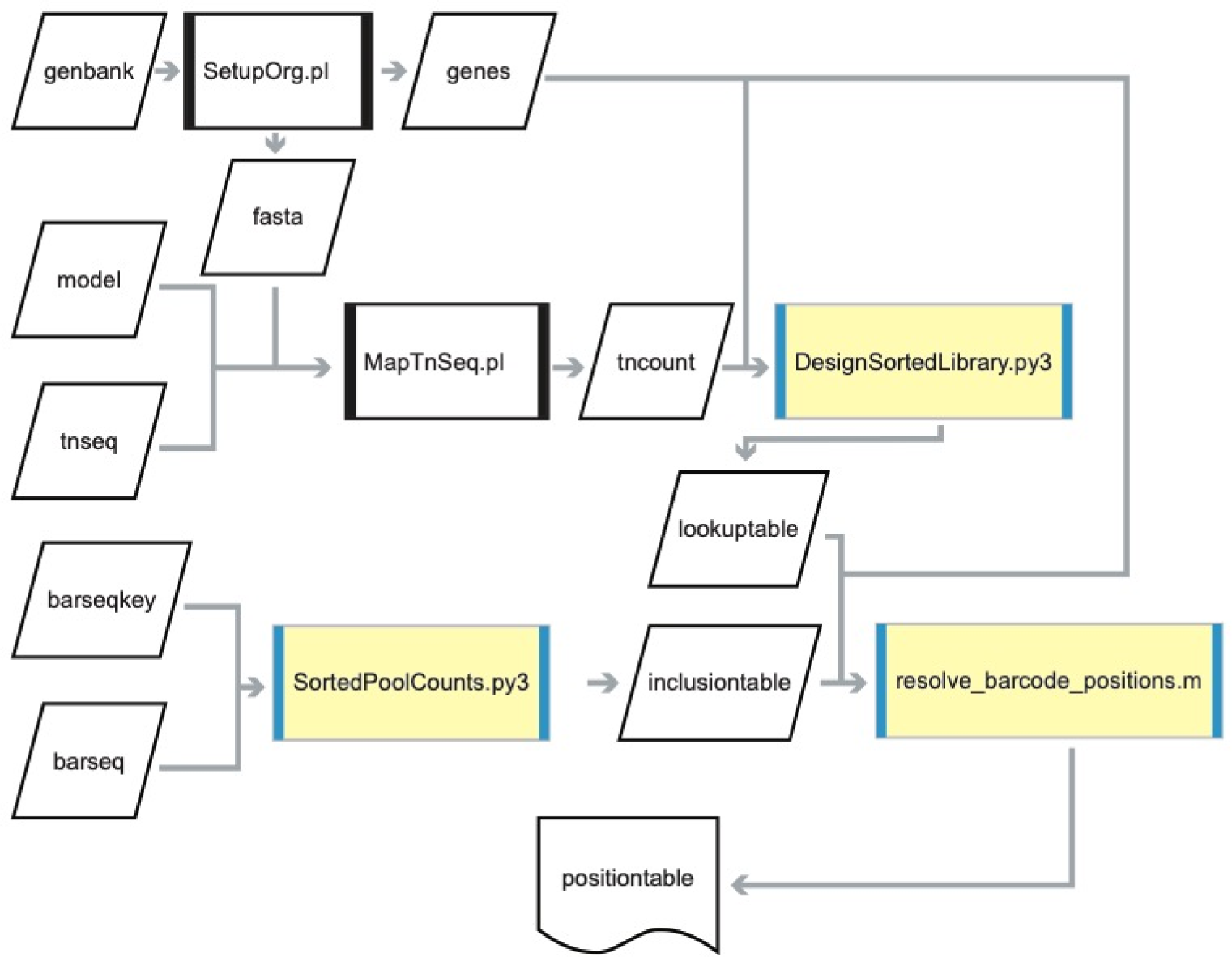
Analysis workflow to locate mutant strains in the ordered library. Files and data are shown as parallelograms. Scripts are shown as rectangles. Rectangles with black edges are scripts from the previously published FEBA code repository. Yellow rectangles with blue edges are scripts written for this study. The language of the scripts is reflected in the file extensions (Perl scripts end in .pl, Python scripts end in .py3, and MATLAB scripts end in .m). See the code availability statement for more information. Input files to the analysis are aligned on the left. The final output, *positiontable*, is at the bottom. SetupOrg.pl takes a genbank-formatted genome sequence (*genbank*) as input as splits it into a simple fasta-formatted sequence (*fasta*) and a table of gene locations (*genes*). MapTnSeq.pl takes the RB-TnSeq data (*tnseq*), a model of the expected transposon sequence in the RB-TnSeq data (*model*), and *fasta* and outputs a table of read counts for every unique combination of barcode and insertion site location (*tncount*). DesignSortedLibrary.py3 takes the information in *tncount* and processes it to create a lookup table connecting unique barcodes to their likely insertion site(s) in the genome (*lookuptable*). SortedPoolCounts.py3 takes the Bar-seq data (*barseq*) and a formatting key (*barseqkey*) and creates a table listing read counts of every barcode in every pool (*inclusiontable*). The final script, resolve_barcode_positions.m, uses the *inclusiontable*, *lookuptable*, and *genes* files to locate transposon insertion strains in the library and record this information in a table (*positiontable*).

### TROUBLESHOOTING

**Table 1:**
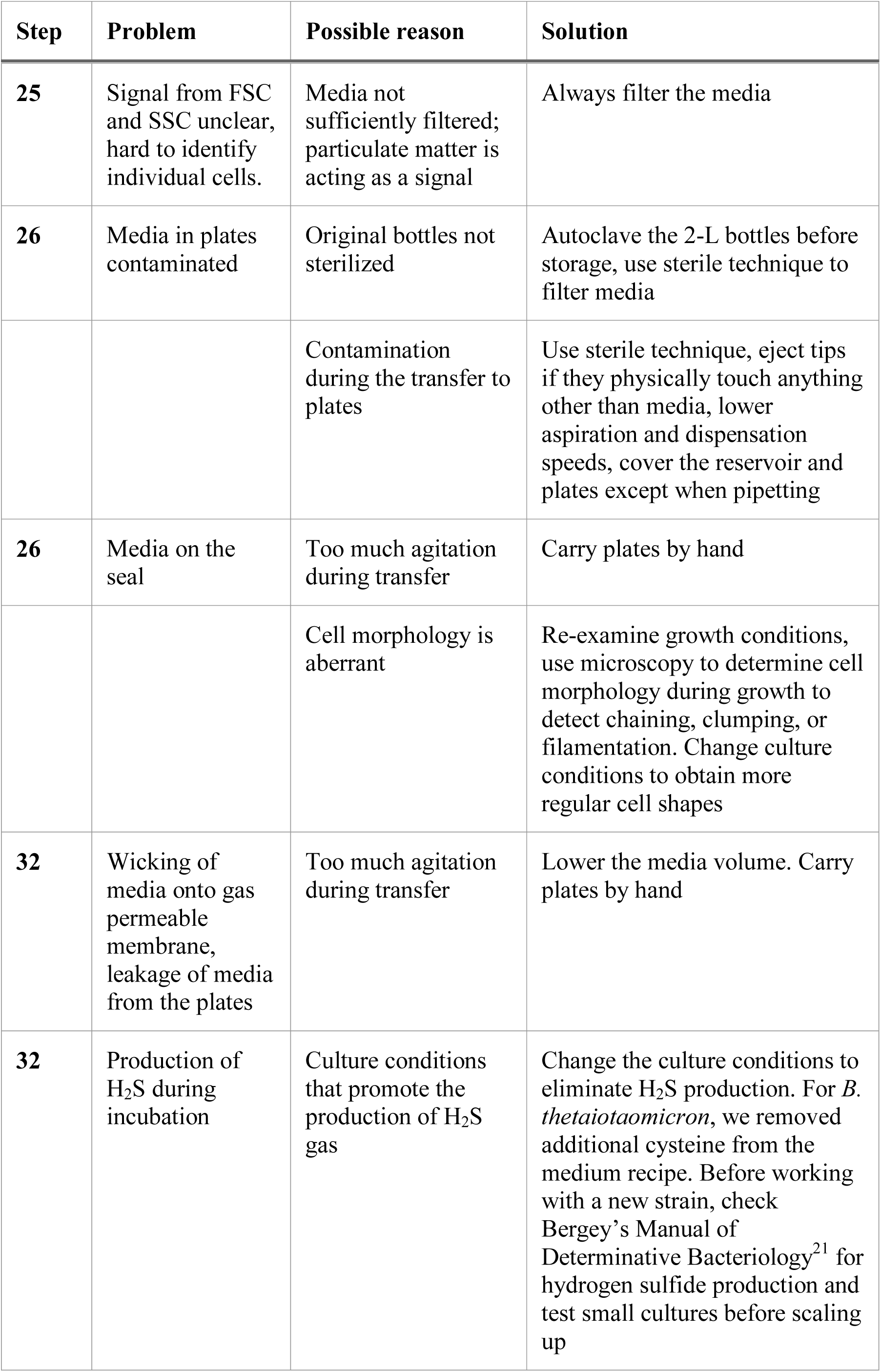
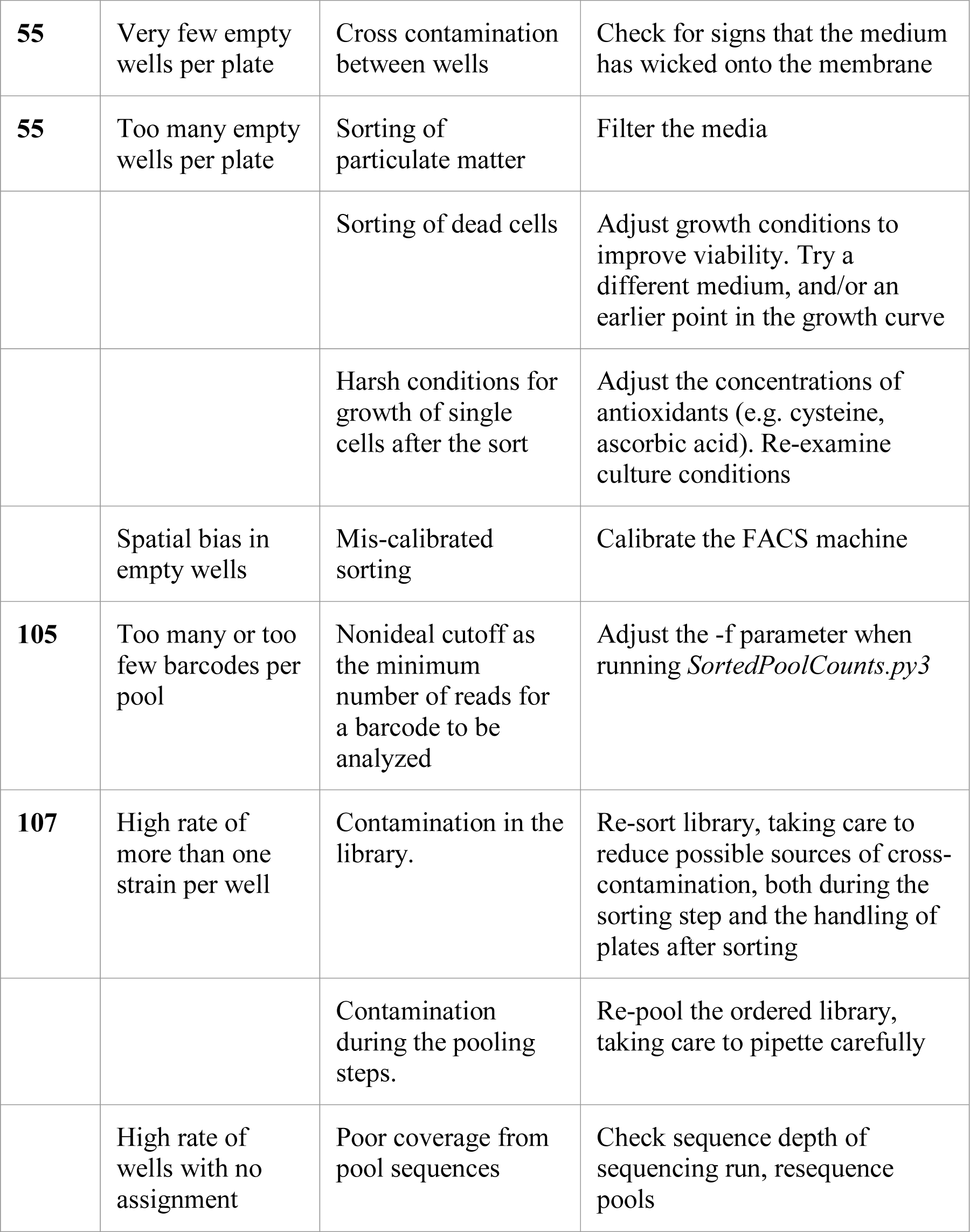
Troubleshooting advice for common problems, with the associated step and possible explanation(s).

### TIMING

Steps 1-11, Fill 96-deepwell plates with medium: 3-4 h for 45 plates

Steps 12-15, Inoculate an overnight culture with a library aliquot: 2 h hands on / 16 h total

Steps 16-18, Grow the library to log phase: 2-5 h (will vary by strain and media)

Steps 19-20, Prepare for the sorting: 0.5 h

Steps 21-31, Sort the library: 2 h for 45 plates

Step 32, Growth of the ordered library: 2-3 d (will vary by strain and media)

Steps 33-35, Labeling of microwell plates: 1 d

Steps 36-37, Transfer of material to anaerobic chamber: 0.5 h

Steps 38-53, Aliquot and store glycerol stocks: 1 d

Step 54, Measure OD_600_ of the library cryostocks: 2 h

Step 55, Analyze cryostock OD_600_ for quality control: 1 h

Steps 56-76, Pool the ordered library for Bar-seq and RB-TnSeq: 1 d

Step 77, Extract genomic DNA from samples: 4 h

Step 78-84, Prepare the Bar-seq library: 1 d

Step 85, Sequence the Bar-seq library: 1 d

Steps 86-94, Prepare the RB-TnSeq library: 1 d

Step 95, Sequence the RB-TnSeq library: 1 d

Steps 96-106, Resolve the positions of barcodes in the library: 3 h

### ANTICIPATED RESULTS

As a test of our approach, we sorted single cells from a barcoded transposon insertion library of *B. thetaiotaomicron* VPI-5482 into 40 96-well plates. We then located barcodes within the ordered library using Bar-seq. We predicted a position for 3517 barcodes within the library, of which 235 were found in more than one position and had to be resolved probabilistically^8^. The final map of transposon insertions in the library identified 3059 wells with a single strain, 350 wells with more than one strain, and 431 wells without a barcode assignment (Figure 7a, top). Less than a third of the wells without a barcode overlapped with wells in which the OD_600_ was less than 0.25, indicating that low cell density is not the main cause of our inability to detect a barcode (Figure 7a, middle). There were no more than three barcodes per well in the library (Figure 7a, bottom). We spot-checked 15 strains (7 single insertions and 8 multiple insertions) using PCR and confirmed that all 15 were correctly located (Supplementary Figure 1, Supplementary Data 3).

**Figure 7:**
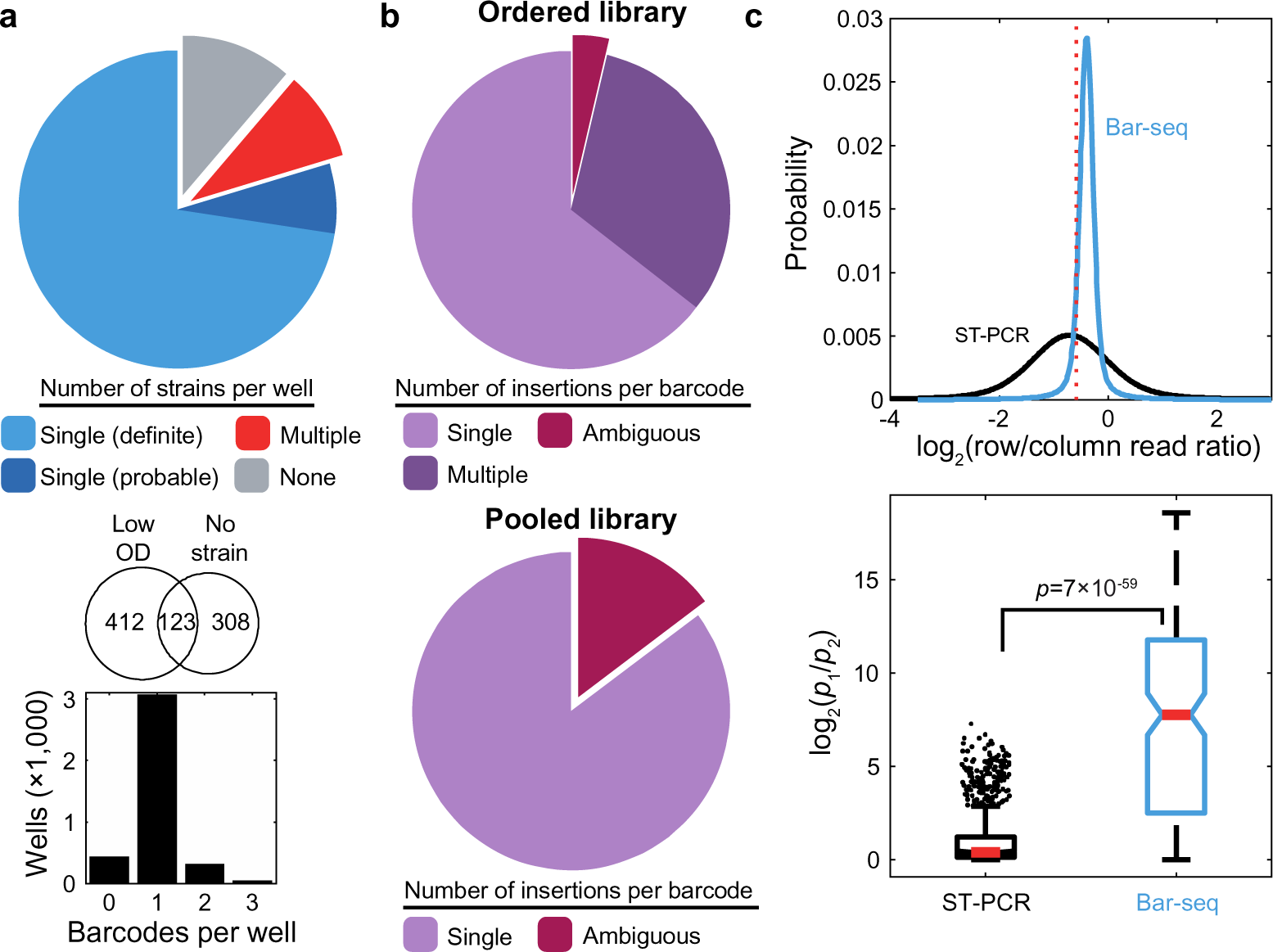
Statistics of strain location identification in a *B. thetaiotaomicron* transposon insertion library sorted into 40 plates. a) Top: we identified 2787 wells carrying a single mutant with a definite solution and 272 wells carrying a single mutant with a probable solution. 350 wells carried more than one strain and 431 wells had none assigned. Middle: only 123 of the 431 wells with no assigned strain also had an OD_600_<0.25, indicating that low cell density was not the sole cause of missing barcodes. Bottom: the majority of wells in the library were assigned a single barcode; no well contained more than 3 unique barcodes. b) Barcodes associated with more than one insertion have an ambiguous assignment in pooled libraries. Performing RB-TnSeq on the ordered library shows that the vast majority of barcodes with more than one insertion are strains with multiple insertions (top), significantly reducing the fraction of ambiguous assignments relative to RB-TnSeq of the pooled library (bottom). c) The predictive power of Bar-seq for probabilistic location of mutants in the library is greater than ST-PCR. Our pool design shares the same row and column pools as a previous ST-PCR study, allowing us to compare the performance of the sequencing methods. Top: the log_2_(ratio of row/column reads) should be a Voigt distribution centered around −0.585^8^ (red dotted line).The fit Voigt distributions for Bar-seq (blue, this study) and ST-PCR (black, our re-analysis of previous work^8^) were normalized to an area under the curve of 1 and plotted. Bottom: The fit Voigt distributions (above) were used to predict the likelihood of row and column solutions for all mutant strains that appeared in exactly two row and two column pools. There are two unique solutions to this 2×2 pool inclusion pattern. The log of the ratio of the probabilities for the more (*p*_1_) and less (*p*_2_) likely solution (log_2_(*p*_1_/*p*_2_)) is plotted for Bar-seq (blue, *n*=169) and ST-PCR (black, *n*=1451). The red dashes are the medians of the distributions, the edges of the boxes are the upper and lower quartiles, whisker length is 1.5 times the inter-quartile range, and the outliers are plotted individually. Bar-seq has a significantly higher log ratio, signifying better predictions.

#### Sorting clarifies otherwise ambiguous transposon insertion events

In the transposon insertion pool used in this study, 85% of barcodes were associated with a single, unambiguous insertion site (Figure 7b, top). As discussed above, the remaining barcodes were ambiguous because they could have arisen either from multiple transposon insertions in the same cell or from separate strains that serendipitously received the same barcode.

It is extremely unlikely that distinct strains with the same barcode would be sorted into the same well, implying that barcodes with a definite location in the library are associated with a single strain. By performing RB-TnSeq on the ordered library as well as on the original collection, we were able to disambiguate multiple barcodes, as barcodes with a definite location that are associated with multiple insertions are likely to be single strains with multiple insertions. Using this logic, we reduced the fraction of ambiguous barcodes in the ordered library to 4% (Figure 7b, bottom).

Notably, the fraction of strains with more than one insertion was greater when predicted from RB-TnSeq of the ordered library compared with the larger pool (Figure 7b). It is possible that decreasing the diversity of the library by more than an order of magnitude allows for greater power in detecting additional insertion sites. Thus, if a substantial fraction of insertions in the pooled library are ambiguous, performing an additional RB-TnSeq reaction on the ordered library may disambiguate many of them.

#### Barcode sequencing improves probabilistic solutions to library positions

In this protocol, we used barcode sequencing (Bar-seq) to sequence pools and locate mutant strains within the library. Alternative sequencing methods that have been used in previous studies include insertion sequencing (INSeq)^5^, transposon sequencing (Tn-seq)^7^, and semi-random two-step PCR (ST-PCR)^8^. Bar-seq and ST-PCR are significantly simpler and more cost-effective than INSeq and Tn-seq; these PCR-based sequencing methods avoid fragmentation, restriction enzyme digestion, size selection, ligation, and multiple purification steps that are required for INSeq and Tn-seq. While ST-PCR is inexpensive and simple, it suffers from inherent noise during the first PCR amplification, in which one of the two primers has semi-random specificity. Because of this issue, we expect that Bar-seq will be more accurate than ST-PCR in quantifying the relative counts of barcodes in the pools, which is important when predicting the probabilistic positions of repeated barcodes in the library.

The pool designs used here and elsewhere^7,8^ are optimized for the time and effort required to pool the libraries. If a barcode occurs in exactly one each of the plate, row, and column pools, then for our pool design it has a definite solution for a single location in the library. If a strain occurs more than once in the library, then its barcode will appear in more than one of each of the different pool types and, with our pool design, there will be multiple possible solutions for its locations in the library. To predict the correct locations of these repeated transposon insertion strains, a previous study^8^ developed a probabilistic solution that we implemented here.

This approach uses the relative read counts of a barcode across pools to predict the most likely solution out of all alternatives. Briefly, the ratio of reads of the same barcode in two pool types (plate, row, or column) should depend only on the number of strains added to each pool. For example, the ratio of reads for a barcode in its row and column pools should be 8/12. When choosing between alternative combinations of pools, if the ratio of reads for one combination is closer to this ideal ratio than the alternatives, then it is the more likely solution. In reality, the ratio of reads is a distribution centered approximately around this ideal ratio due to both systematic and random noise introduced during the amplification process. A tighter distribution will be more predictive when choosing alternatives.

Because our pool design (plate, row, column) and the pool design from a previous study using ST-PCR (plate-row, plate-column, row, column)^8^ share the row and column pool types, we were able to directly compare Bar-seq to ST-PCR according to their ability to distinguish between alternative solutions for indefinite strain locations. We first examined the distribution of row versus column read ratios for definite solutions in the two libraries. The log_2_(row/column read ratio) should be a Voigt distribution centered around −0.585^8^. We found that the Bar-seq method (this study) resulted in a tighter distribution than ST-PCR (reanalyzed data from previous work^8^) (Figure 7c, top), indicating that Bar-seq better preserved read count information contained in the genomic DNA pools.

We next examined the subset of barcodes (this study, *n*=169) or transposon insertion locations (reanalyzed data from previous work^8^, *n*=1451) that were detected in exactly two row and two column pools. There are two unique solutions that could give rise to this pool inclusion pattern. The log ratio of the probabilities of the more (*p*_1_) and less (*p*_2_) likely solutions is a measure of the ability of read ratio information to predict a correct solution for the mutants. The Bar-seq method outperformed ST-PCR (Wilcoxon rank sum test: *W*=2×10^5^, *p*=7×10^−59^) (Figure 7c, bottom).

We conclude that Bar-seq is more robust than ST-PCR and more cost-effective than INSeq and Tn-seq, making it the preferred sequencing method for locating mutants within an ordered library.

#### Limited growth in certain wells is likely due to transient oxygen stress during sorting

After aliquoting and storing the library, we measured the OD_600_ after ∼60 h of growth. In 14% of the wells, the OD_600_ fell below a threshold of 0.25, but in many cases was significantly higher than the expected OD_600_ of a blank well (Figure 8a). Microscopic examination of the cultures in these low OD_600_ wells revealed that the majority still had detectable culture densities (Figure 8a,b). Furthermore, the cultures in these low OD_600_ wells could be recovered by outgrowth (Figure 8c), with a longer lag time as the main feature that distinguished them from wells that exhibited high OD_600_ values during initial outgrowth (Figure 8d). To test if the low OD_600_ was connected to a biological process disrupted by the transposon insertions, we looked for a correlation between low OD_600_ and the disrupted genes of the transposon insertion strains. Of the 61 genes that had more than one insertion strain isolated as a pure culture in the ordered library, we found that one strain having a low OD_600_ did not make it more likely that the second also had a low OD_600_. In fact, there was zero overlap between pairwise comparisons (Fisher’s exact test: *p*=0.6, odds ratio=0). We conclude that low OD_600_ wells in the *B. thetaiotaomicron* ordered library are not due to a fitness defect of the mutant that was sorted. Instead, we hypothesize that the transient exposure to oxygen during sorting temporarily slows growth for a fraction of the sorted wells. The presence of these low OD_600_ wells could be a useful indication that post-sort recovery conditions of a strain can be further optimized.

**Figure 8:**
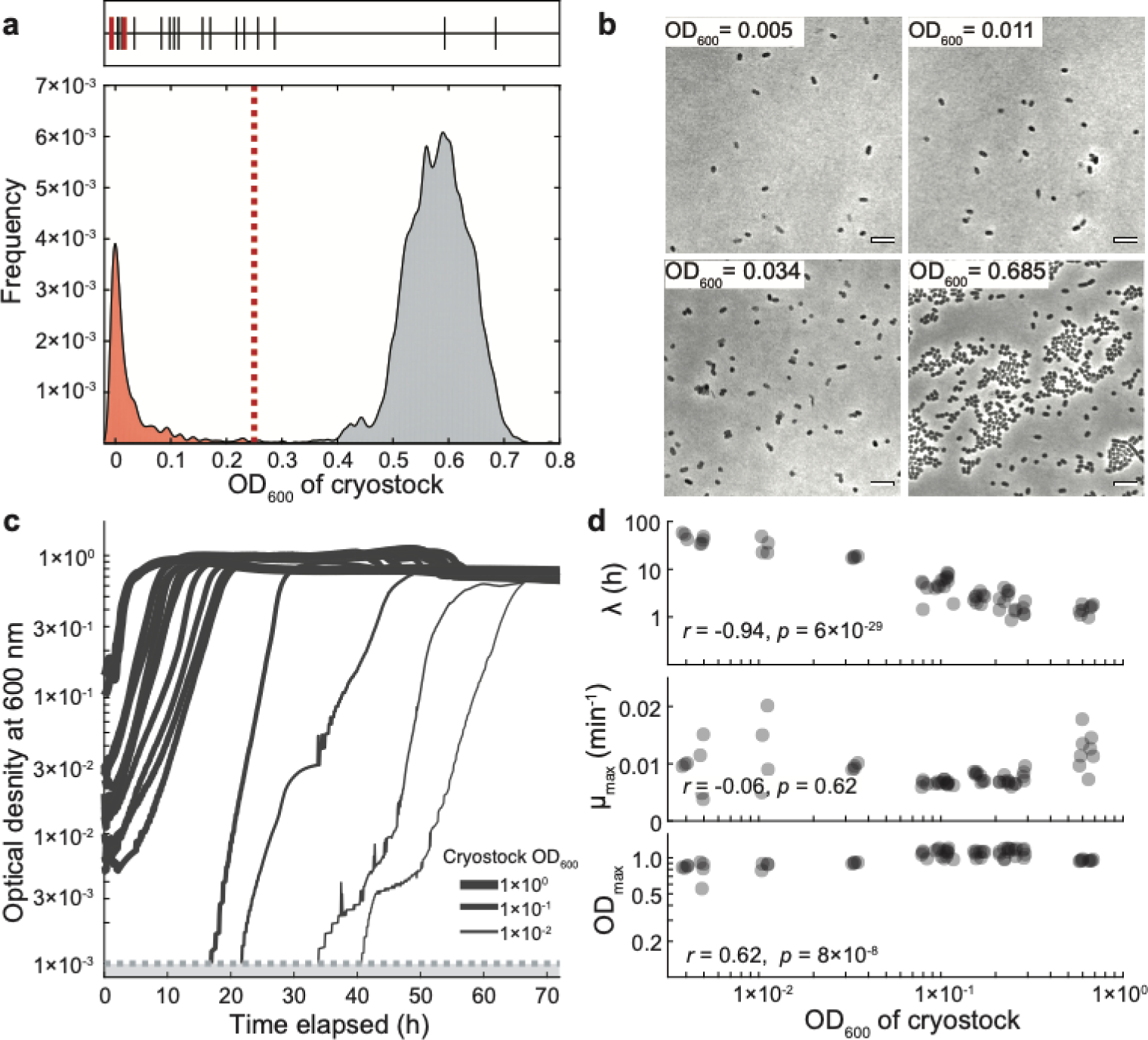
Growth properties of the 40-plate *B. thetaiotaomicron* barcoded transposon insertion library. a) The distribution of OD_600_ measurements in the cryostocks of the ordered library revealed a surprisingly large population of low OD_600_ wells. A heuristic cutoff of 0.25 (red dotted line) was chosen to discriminate between high (grey distribution) and low OD_600_ (red distribution) in the library. Tick marks at the top of the distribution denote the OD_600_ of wells that were chosen for investigation. Black tick marks record wells in which cells were found by microscopy, while the two red tick marks represent the OD_600_ of wells in which no cells could be found in a 1-µL aliquot. b) Cells were found using microscopy in most of the low OD_600_ wells in the library, even in wells with OD_600_<0.01, indicating that many of the low OD_600_ wells are not due to a problem with sorting. Scale bar: 5 µm. The cryostock OD_600_ in the library is shown for each image/well. Images were acquired with a Nikon Ti-E outfitted with a 100X (NA 1.4) objective and an Andor Zyla 5.5 sCMOS camera. Micromanager was used for image acquisition^25^. c) Growth curves after recovery from the cryo-stock showed that many of the low OD_600_ wells can recover to wild-type densities. The thickness of the line is proportional to the log of the cryostock OD_600_. All curves represent the mean of (*n*=4) independent measurements. The horizontal line at an OD_600_ of 0.001 represents the detection limit of the plate reader. The two wells in which cells were not detected did not recover during the 72 h period of monitoring and are not plotted. d) Growth parameters extracted from the recovery curves (*n*=4) show that the transposon insertion strains isolated from low OD_600_ wells have maximum OD_600_ values (OD_max_) and maximum growth rates (μ_max_) similar to wild-type. As expected, the lag phase (*λ*) of the cultures was negatively correlated with the final OD_600_ in the library (Pearson’s *r*=-0.94, *p*=6×10^−29^).

#### Adaptations to this protocol

This protocol introduces two new advances in creating ordered mutant libraries: flow sorting to isolate single cells of anaerobic species, and barcode sequencing to locate mutant strains within the ordered library.

Sorting using a FACS machine is a fast method for isolating mutants from a large pool that is able to sort multiple anaerobic species when used in combination with an antioxidant (Figure 2). However, it may not be the best approach in all situations. Certain strains may be too sensitive to oxygen to survive sorting and will need to remain in an anaerobic environment for the entire protocol. If the number of strains in the original pool is not significantly greater than the number of strains to be sorted, the same mutant strains are likely to be isolated more than once in the ordered library. Finally, a core facility with FACS machines may not be available. In these cases, it may be preferable to directly pick transformed colonies from a selective plate. Automated colony picking equipment with a footprint small enough to install in an anaerobic chamber is commercially available^23^. Regardless of the approach used to order the library, barcode sequencing still promises to greatly decrease the cost and effort of locating mutations in the ordered library.

Our strategy was to pool according to plates, rows, and columns (Figure 1). This pool design scales linearly with the number of plates in the library, where *N*_pools_ = *N*_plates_ + 20. For larger libraries, previous approaches have used an alternative pool design (plate-row, plate-column, row, column)^7,8^ that scales with the square root of plate number (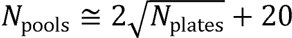). Our plate-row-column pool design is less sensitive to ambiguity from multiple occurrences of the same strain in the library, making it easier to probabilistically locate these strains. For example, a strain that is isolated twice in the library will have 4 potential solutions in a plate-row-column pool design compared with 8 potential solutions in the alternative four-dimensional pool design. Furthermore, the ease of Bar-seq reduces the relative cost savings of a pool design that scales with the square root of plate number compared to linearly. Nonetheless, ordered libraries with a larger number of plates may benefit from this alternative pool design. Similarly, ordered libraries with a high degree of repetition (same mutant, multiple locations) would benefit from a DNA Sudoku pool design^17^. Regardless of the pool design used, we expect that Bar-seq will improve the quality (Figure 7c) and reduce the cost and effort to sequence pools. Small modifications to the tables *lookuptable* and *inclusiontable* that are produced at the beginning of our analysis pipeline should be sufficient for adapting Bar-seq to analysis of alternative pooling designs.

In conclusion, we have demonstrated a single-cell sorting protocol coupled to barcode sequencing that can rapidly and straightforwardly generate a sorted mutant library for anaerobic species. By lowering the investment required to generate an ordered library of mutants, we expect that this protocol will be a powerful advance for researchers investigating genotype-phenotype relationships in the human microbiota, catalyzing new approaches to study the effect of the microbiota on health and disease.

## Data availability

The sequencing data used in this study are publicly available on NCBI as part of BioProject PRJNA573294. The sequence reads from the Bar-seq experiment on pools of the ordered library are available as part of BioSample SAMN12807646. The sequence reads from the RB-TnSeq experiment on the ordered library are available as part of BioSample SAMN12809978. Plate reader output files, microscopy images, and extracted growth parameters that support the findings of this study (Fig. 2, 3, 6) are available from the corresponding author (K.C.H.) upon request.

## Code availability

The code required for locating insertion mutants in ordered libraries is available on BitBucket (https://bitbucket.org/kchuanglab/resolve_barcode_position/src). These scripts rely on previously published code available on BitBucket (https://bitbucket.org/berkeleylab/feba/src).

## Author Contributions

A.L.S., A.M.D., and K.C.H. conceived the study. A.L.S. designed experiments and collected data. A.L.S. and R.C. analyzed the data. A.L.S., R.C., A.M.D., and K.C.H. interpreted the data and wrote the manuscript.

## Acknowledgements

The authors thank the Huang lab for helpful discussions. This work was supported in part by the Allen Discovery Center at Stanford University on Systems Modeling of Infection (to K.C.H.). K.C.H. is a Chan Zuckerberg Biohub Investigator. The authors also acknowledge the hospitality of the Aspen Center for Physics, which is supported by NSF grant PHY-1607611. The following reagents were obtained through BEI Resources, NIAID, NIH as part of the Human Microbiome Project: *Lactobacillus reuteri*, strain CF48-3A, HM-102; *Clostridium innocuum*, strain 6_1_30, HM-173; *Clostridium symbiosum*, strain WAL-14163, HM-309; *Bacteroides finegoldii*, strain CL09T03C10, HM-727; *Parabacteroides johnsonii*, strain CL02T12C29, HM-731.

## Supplemental Information

**Supplementary Data 1:**
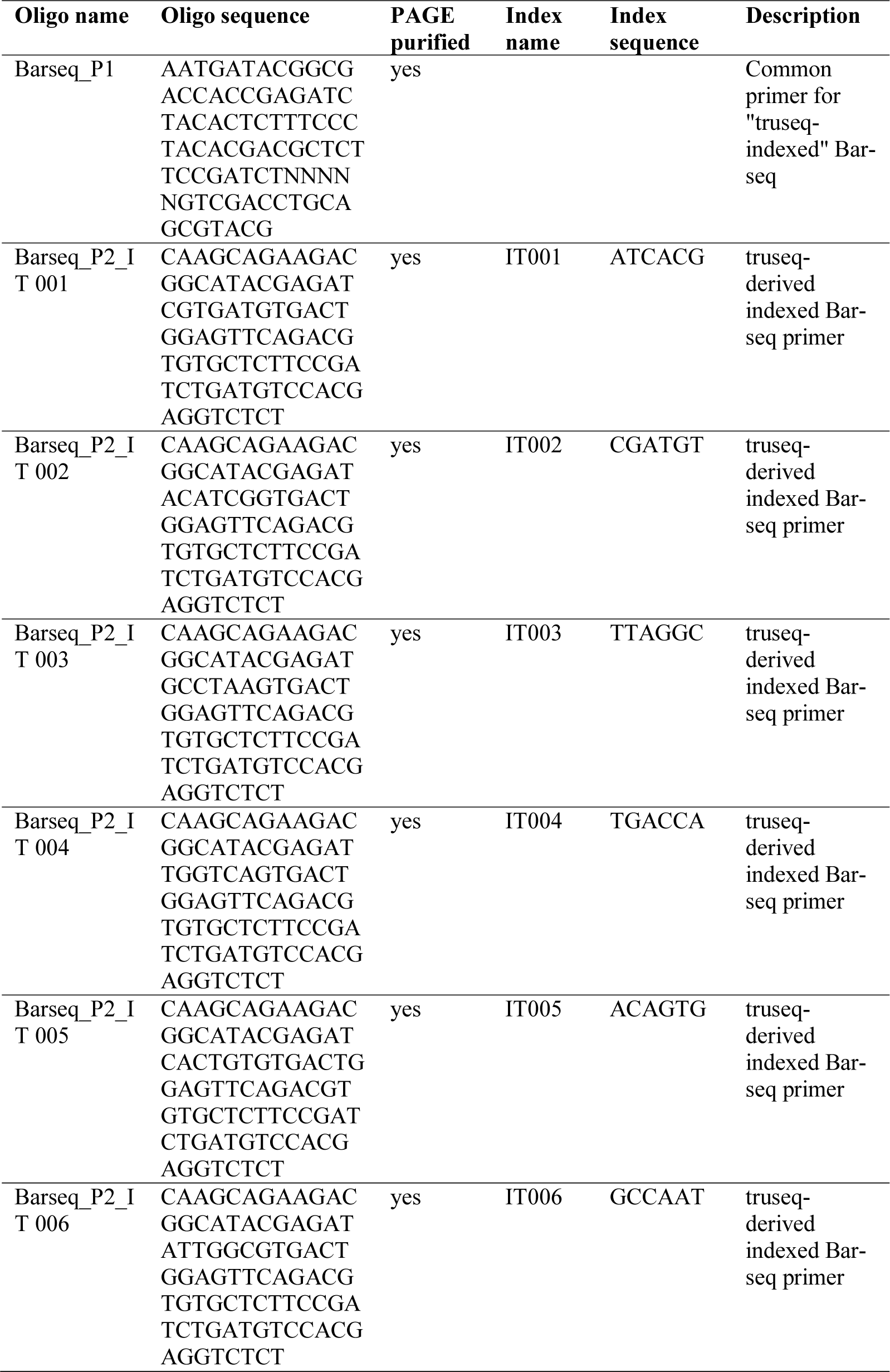
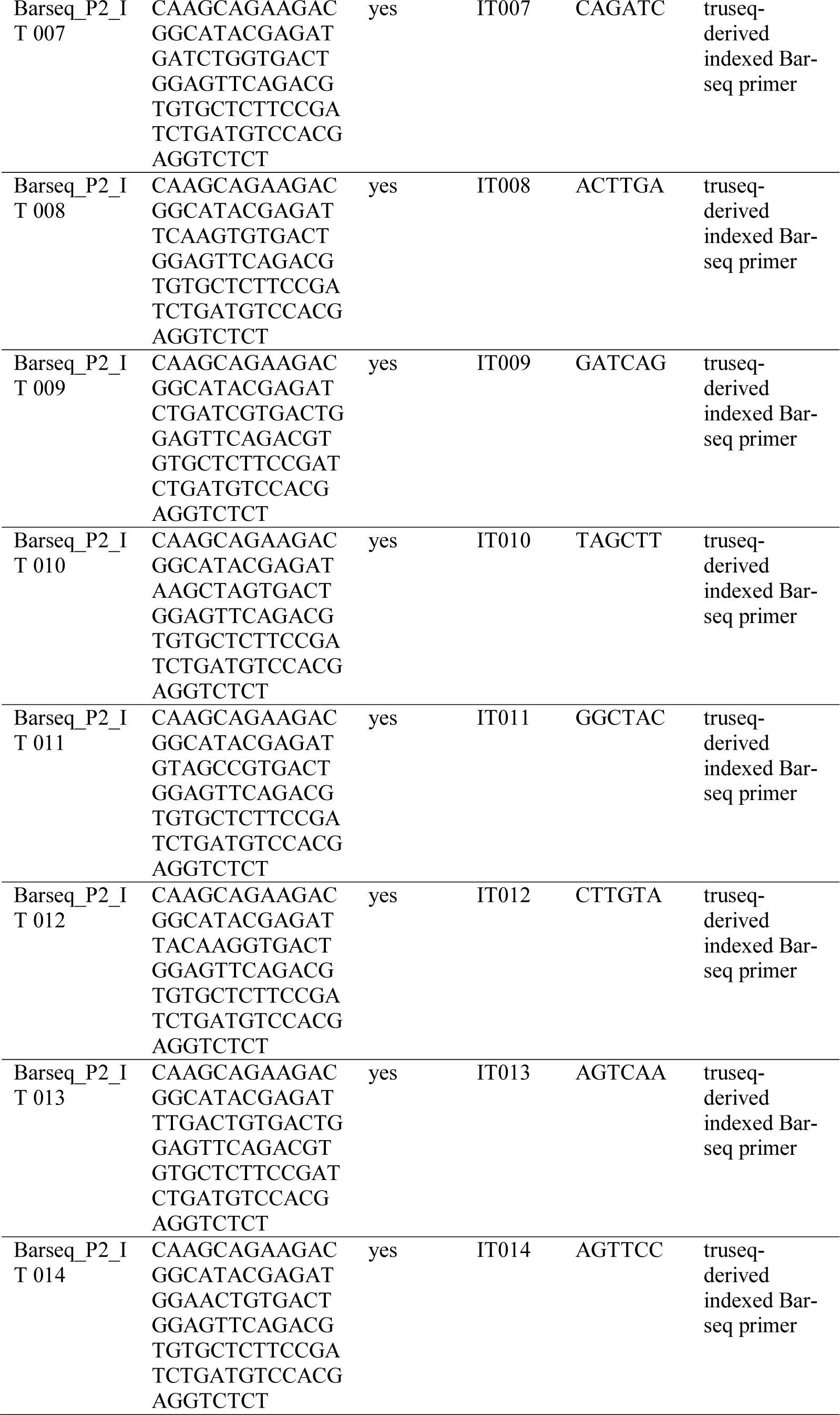
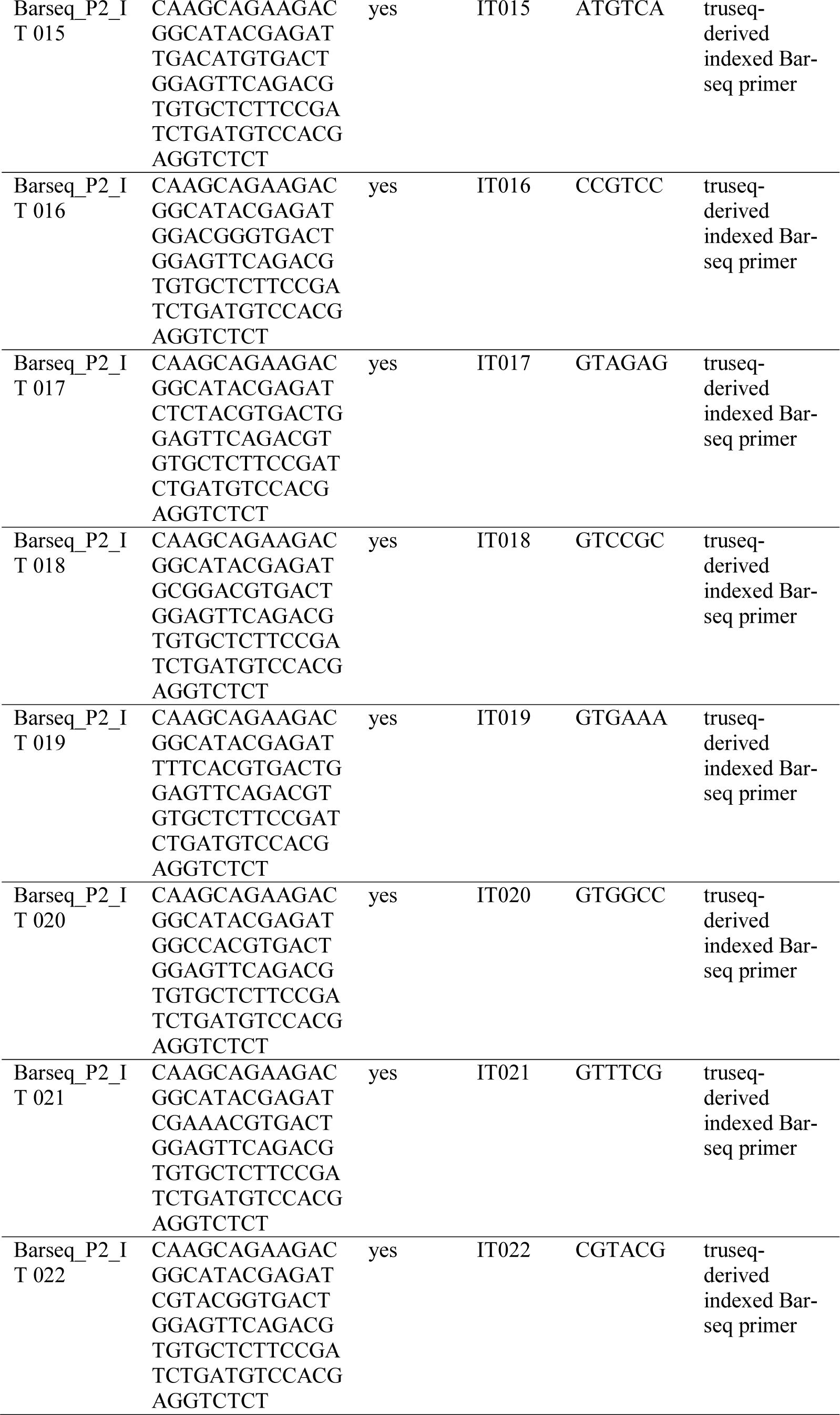
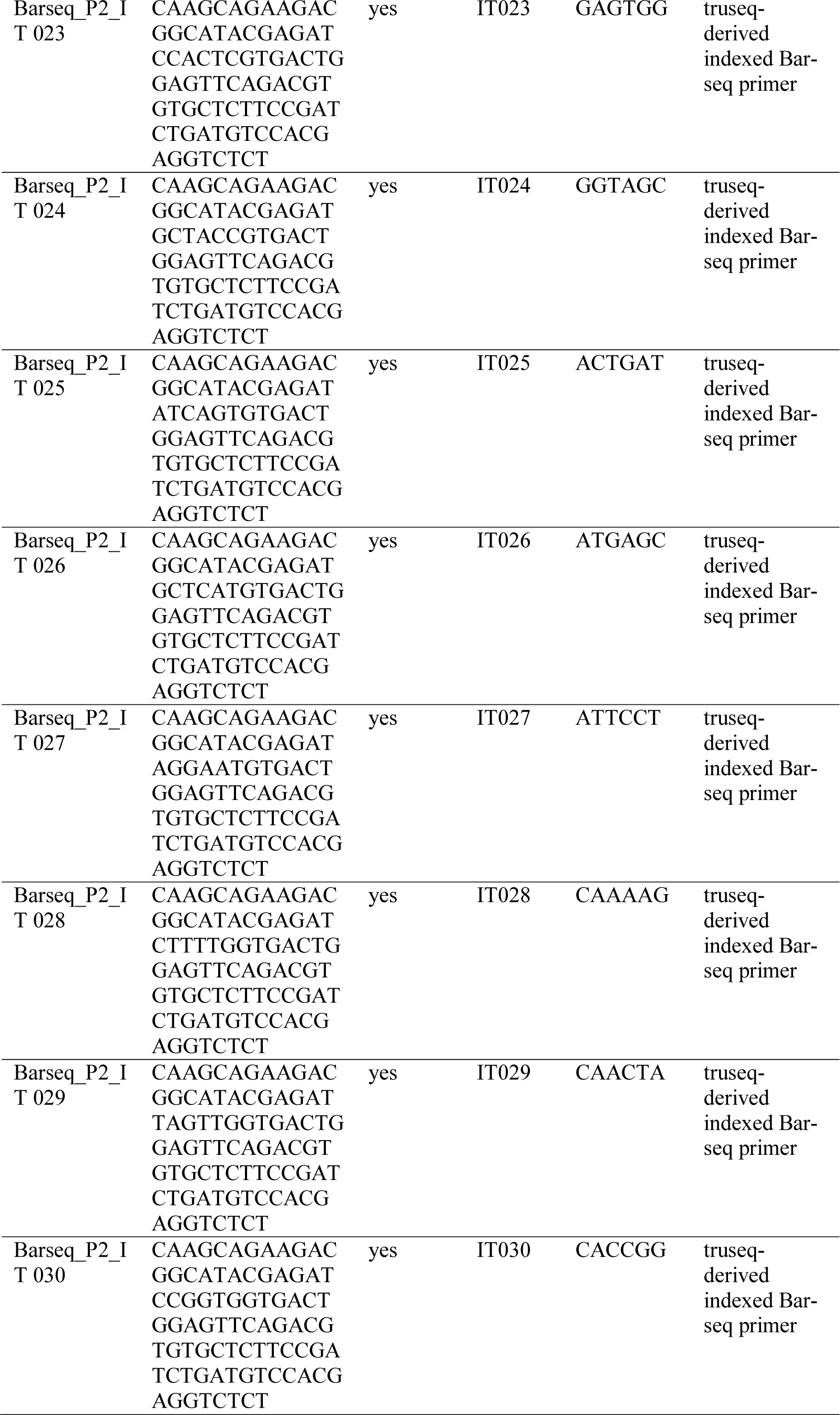
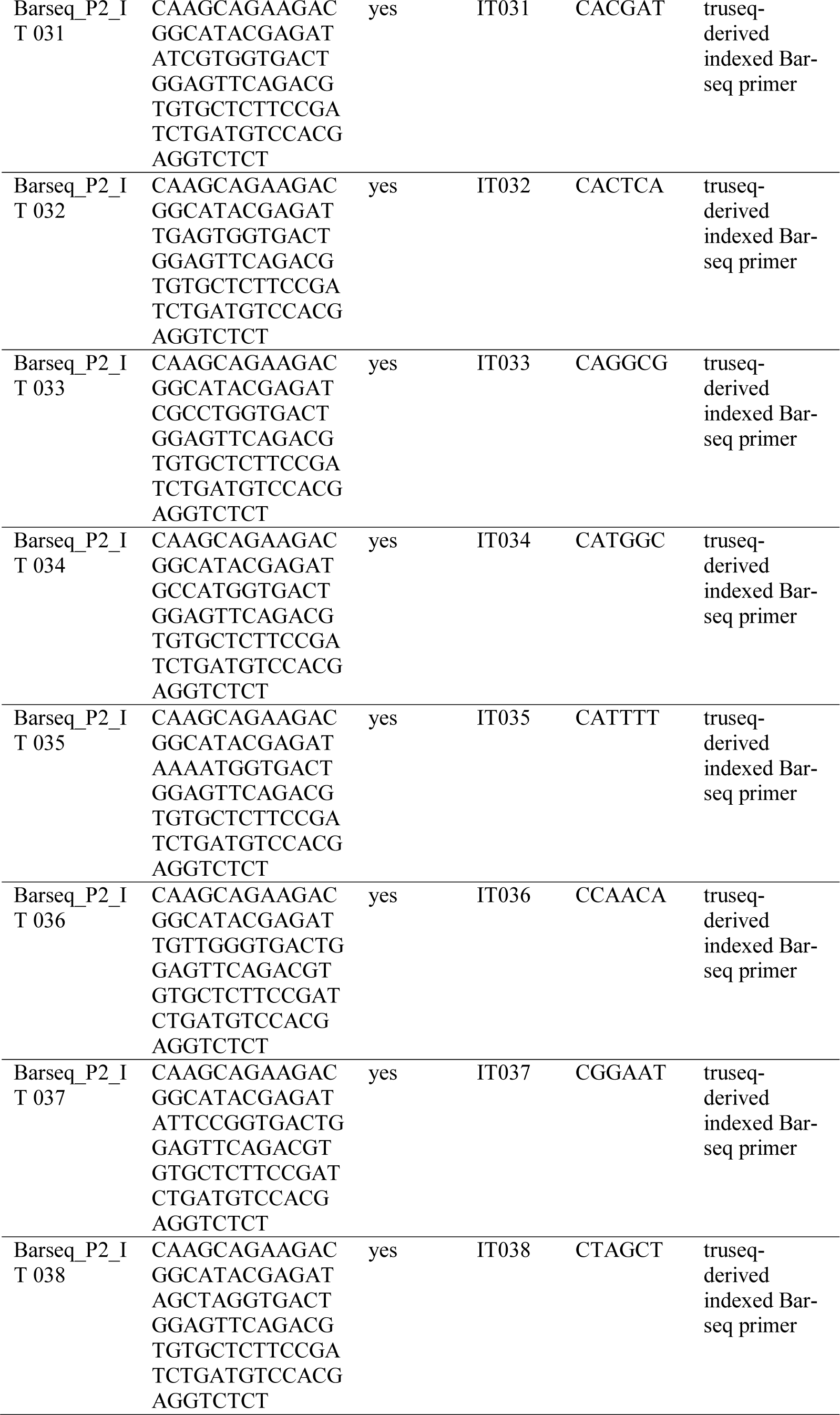
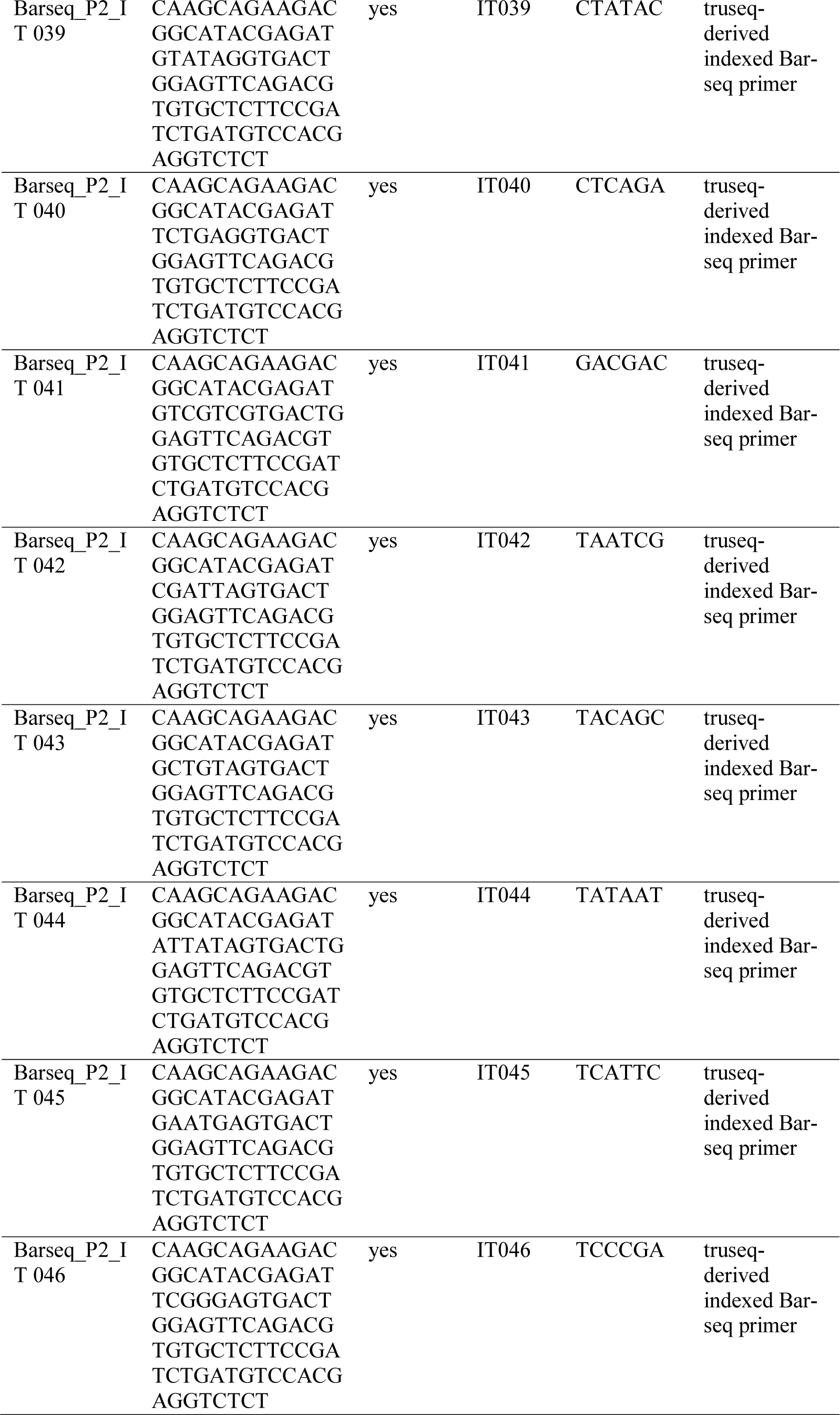
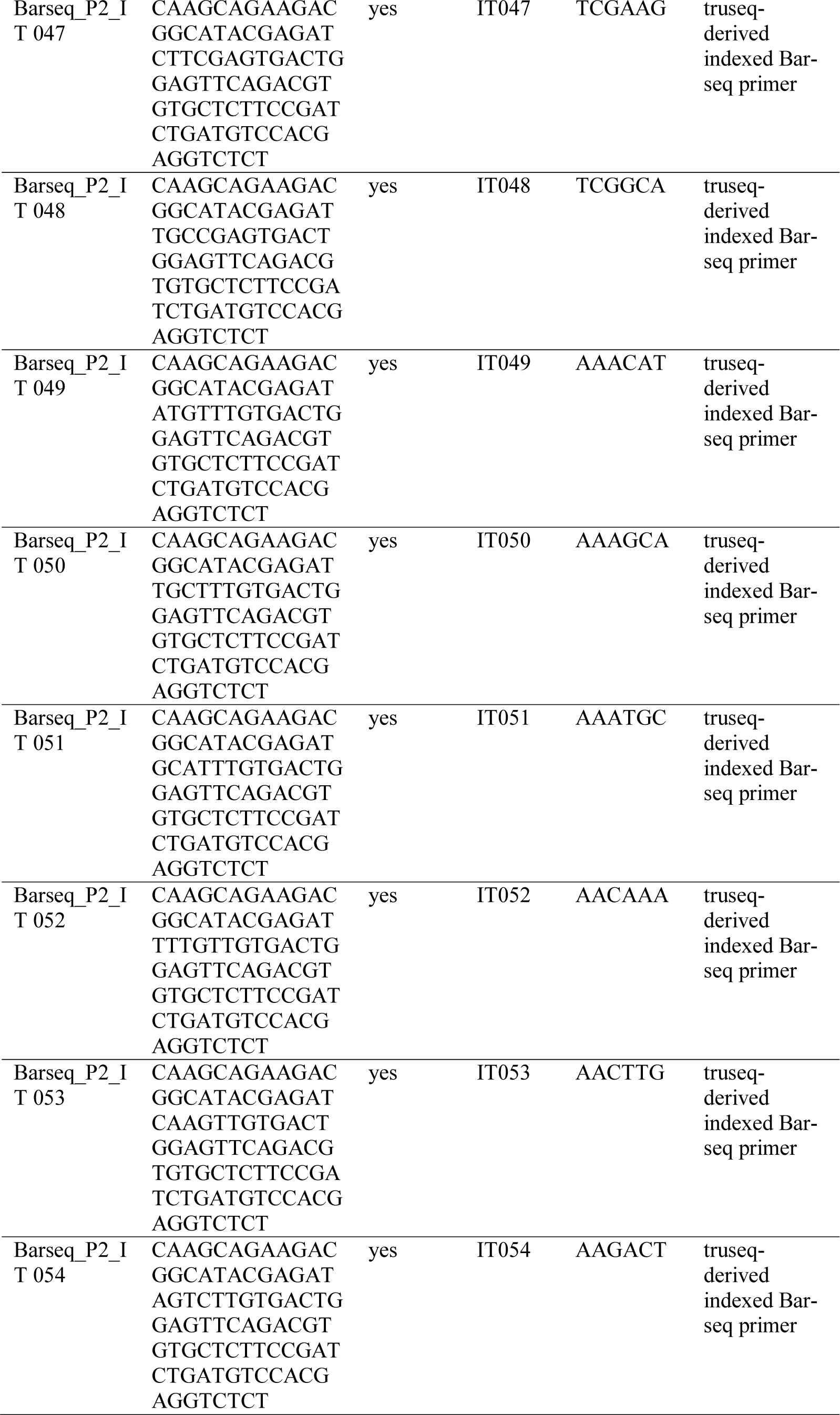
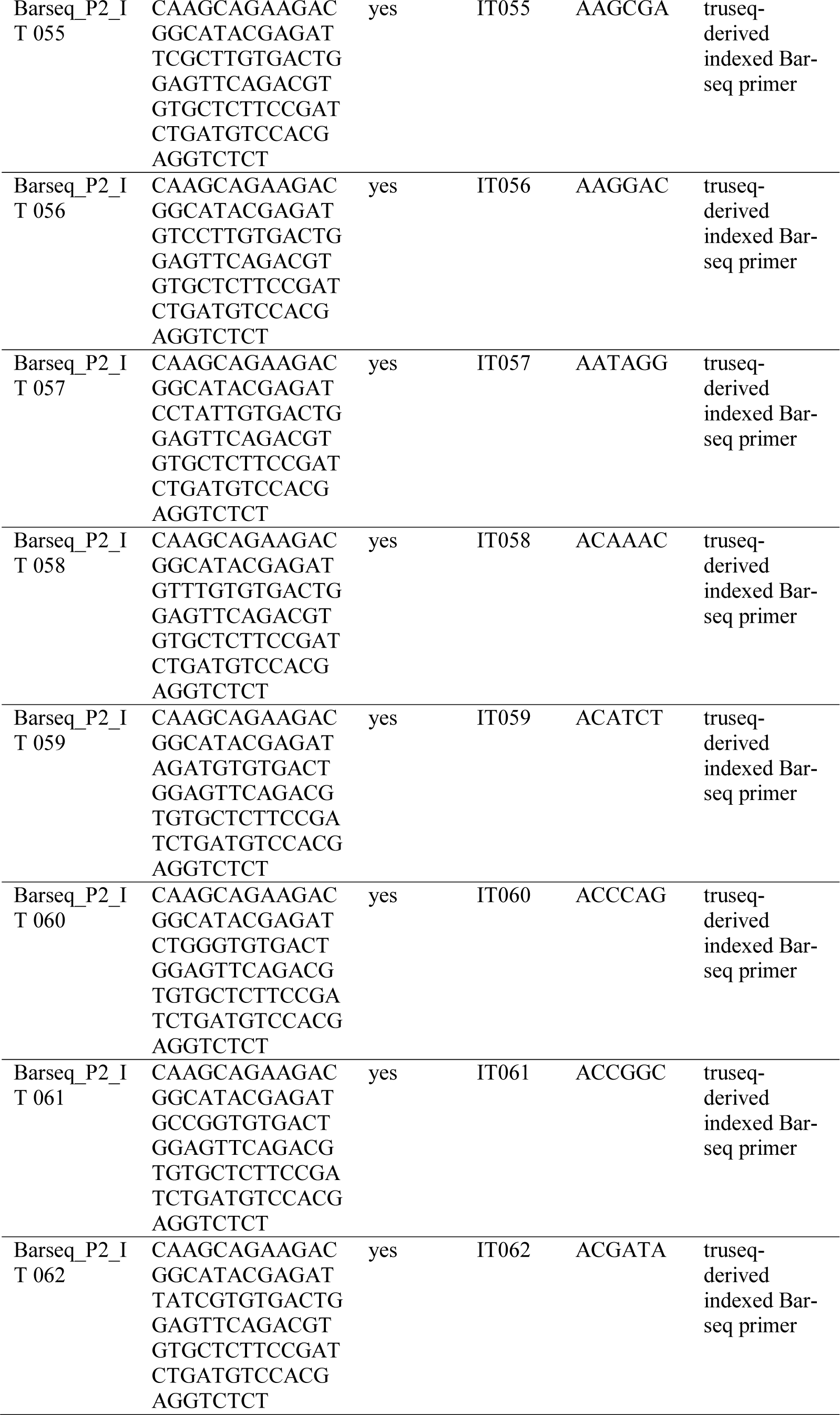
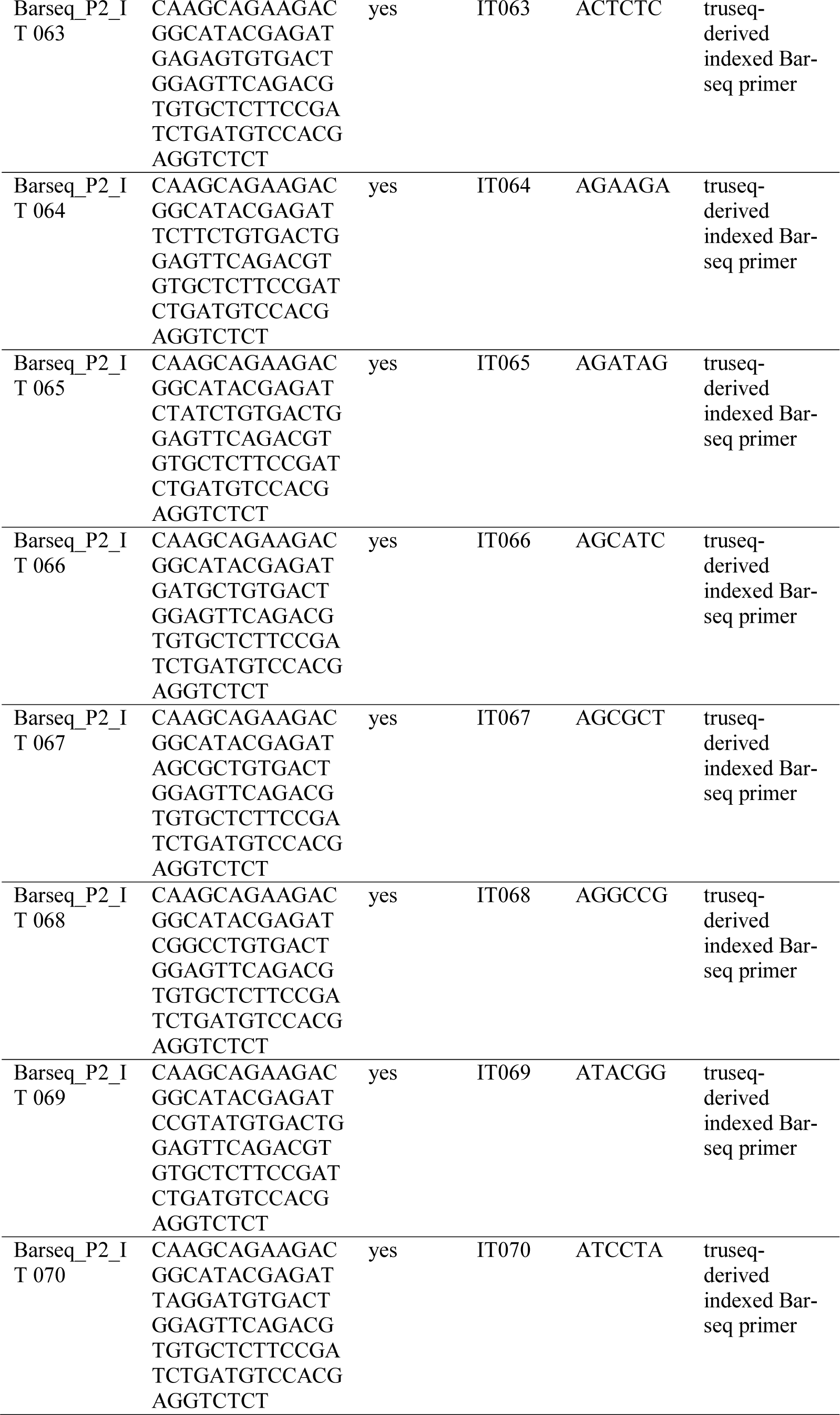
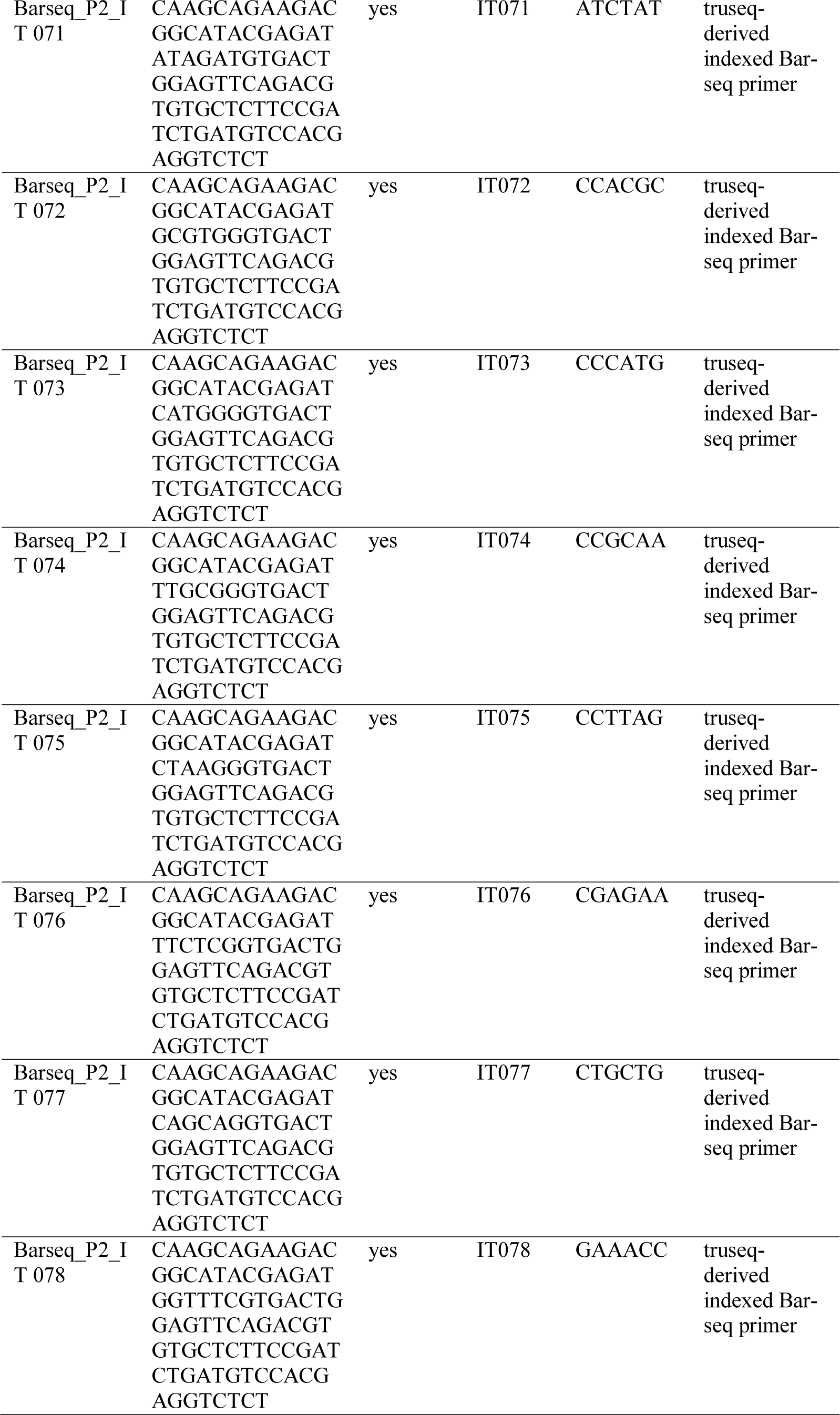
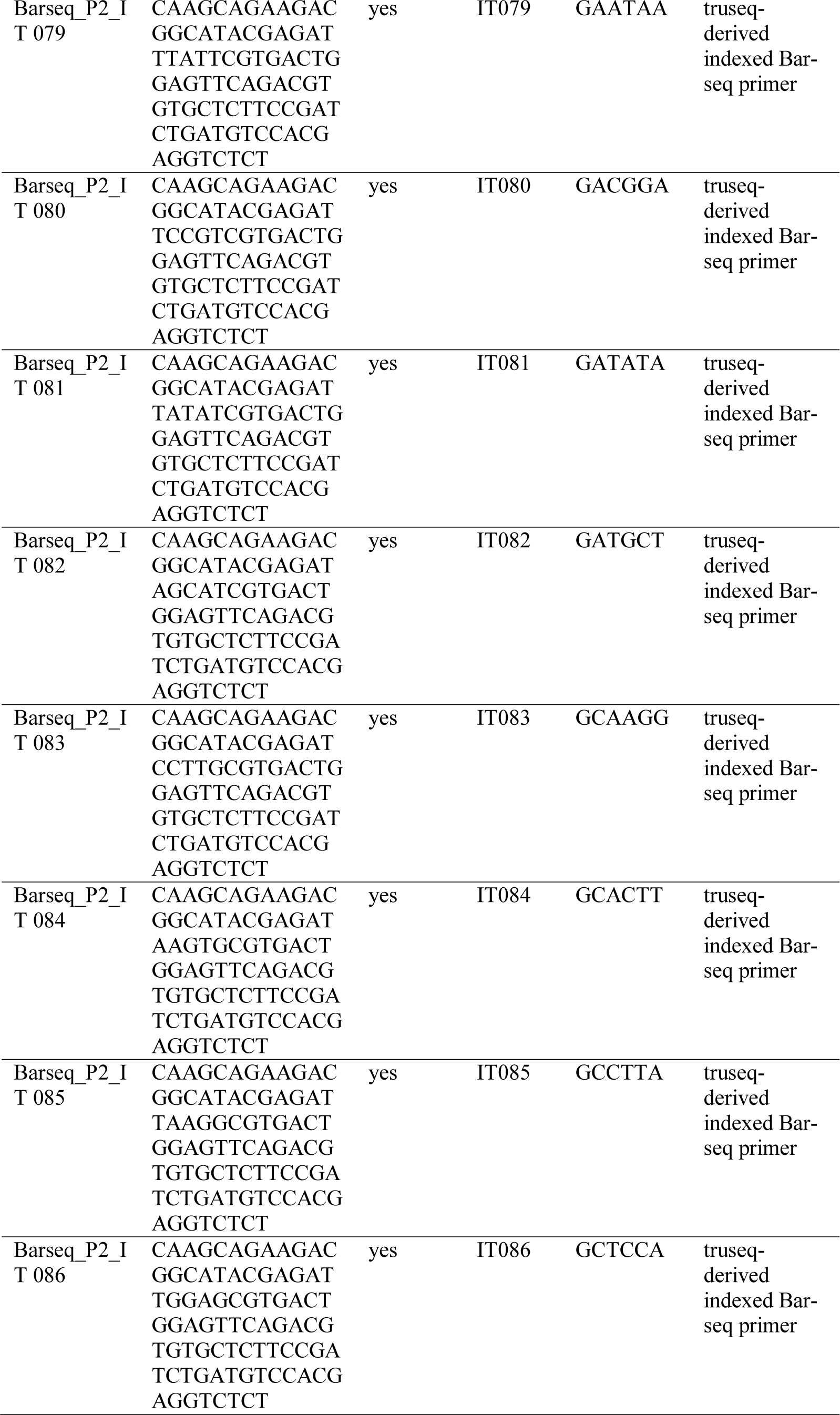
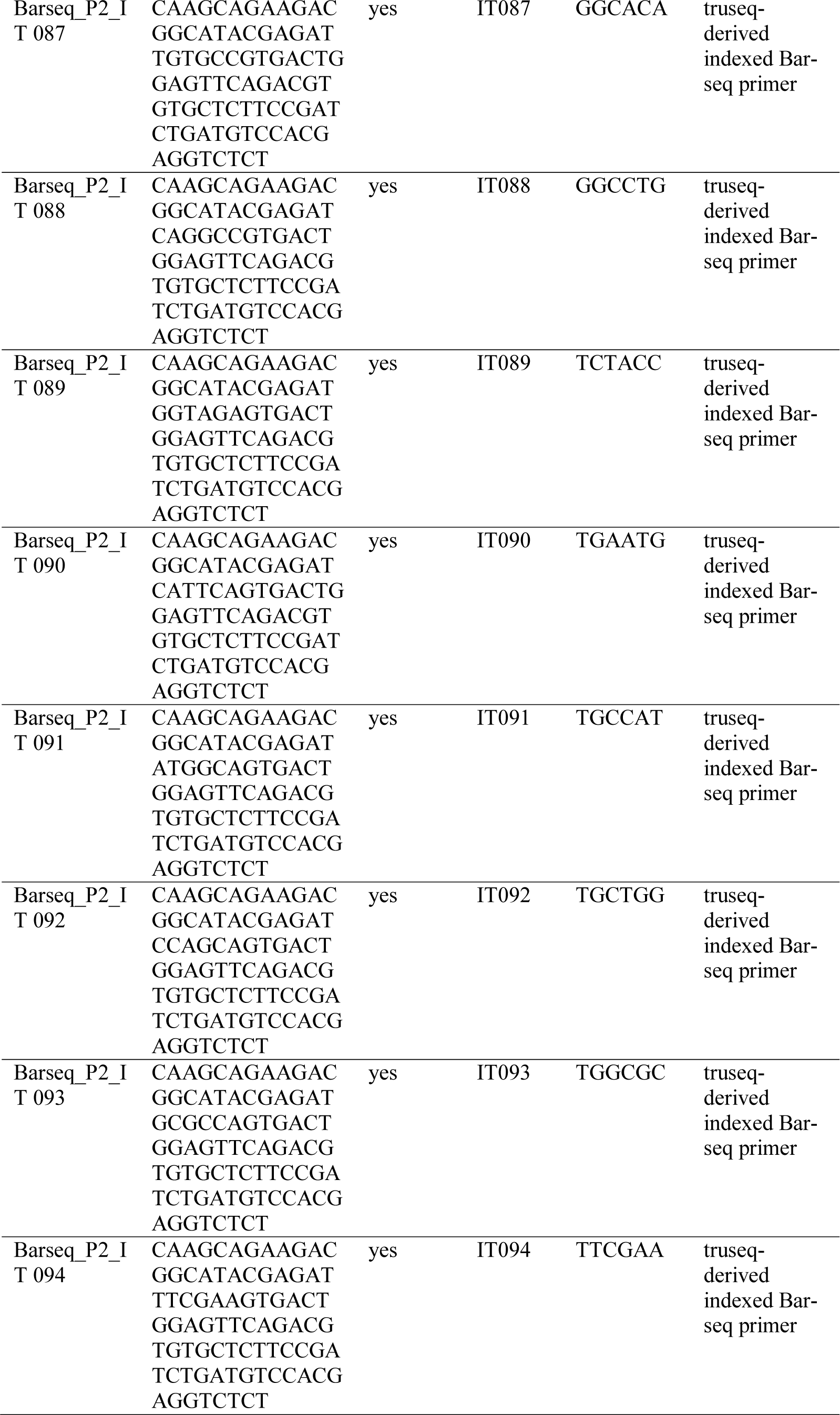
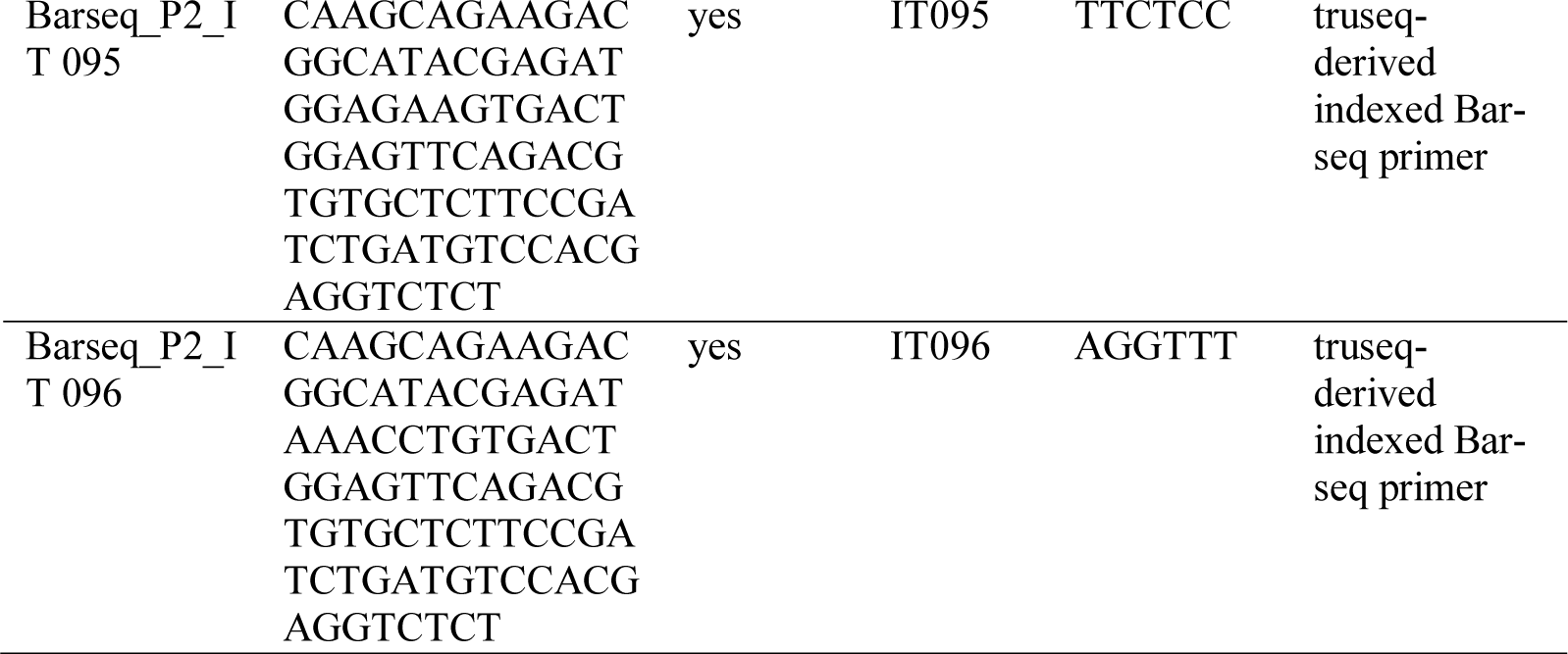
Table of oligos used for Bar-seq.

**Supplementary Data 2:**
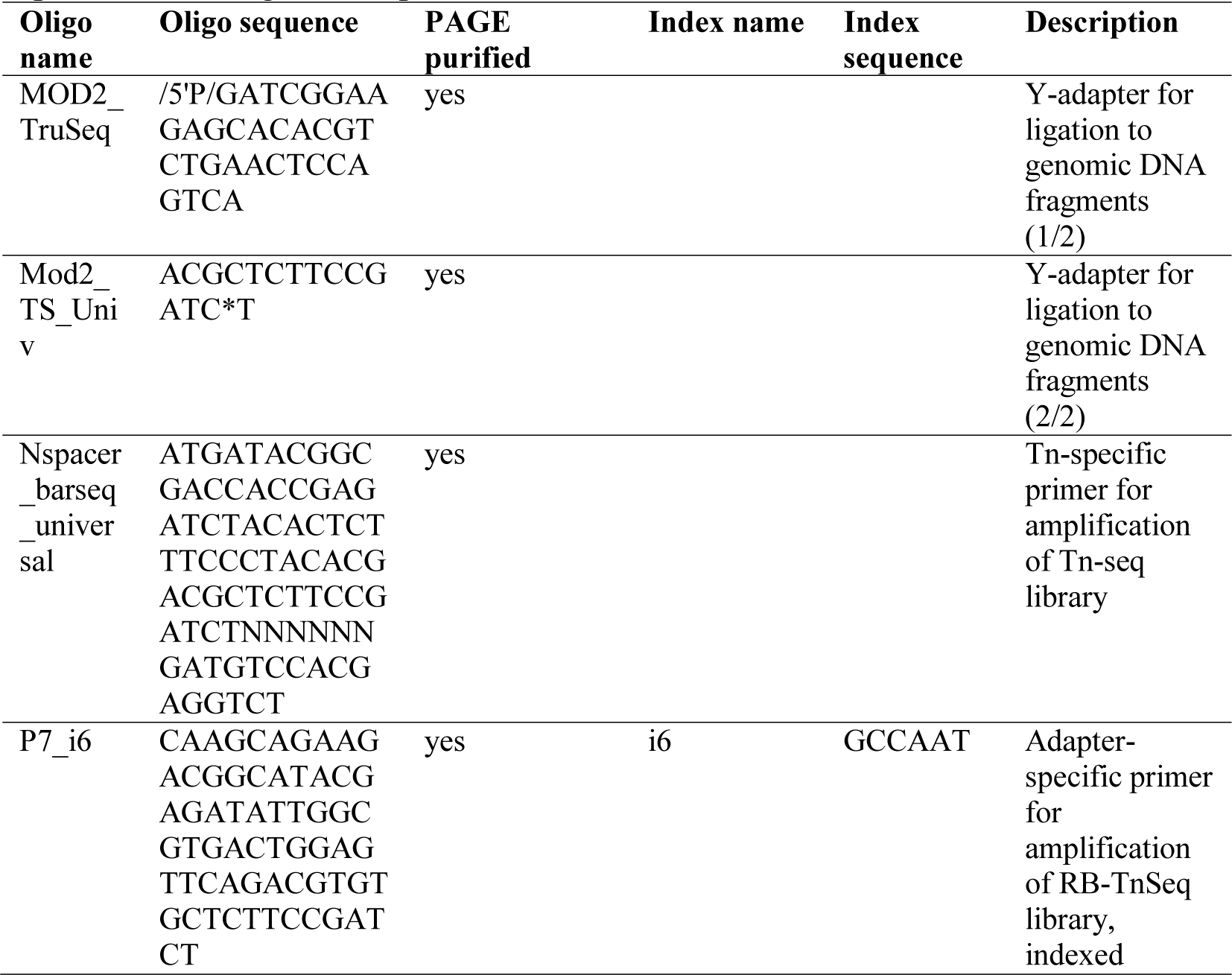
Table of oligos used for RB-TnSeq. The oligo MOD2_TruSeq is modified to carry 5’-phosphate, as denoted by ‘/5’P/’. The oligo Mod2_TS_Univ is modified by an internal phosphorothioate bond, as indicated by the ‘*’. The MOD2_TruSeq and Mod2_TS_Univ oligos are annealed to create a Y-adapter, as specified in Reagent Setup.

**Supplementary Data 3:**
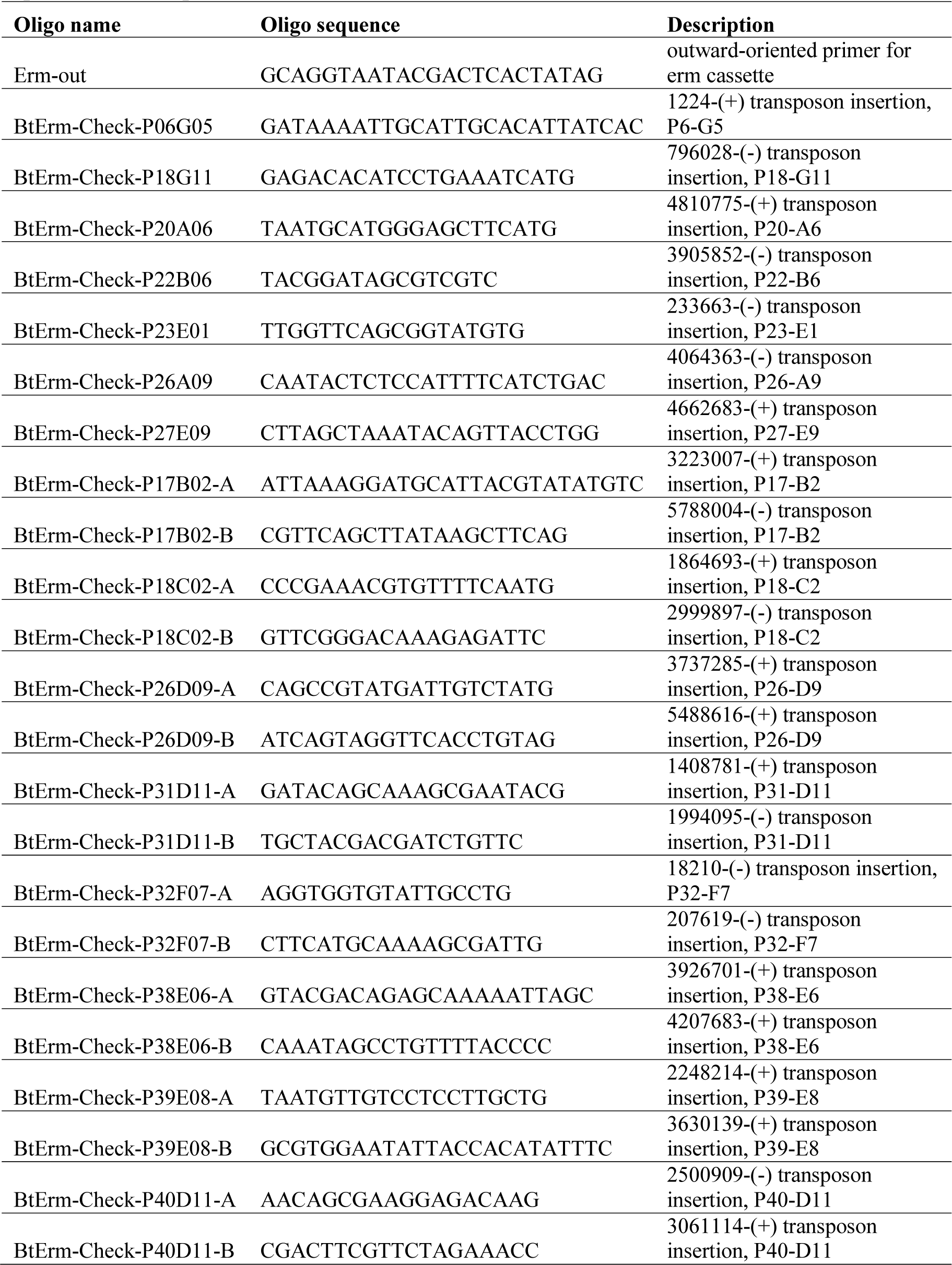
Table of oligos used to spot check strain assignments in the ordered library. The transposon insertion targeted for the PCR check is annotated with position-(orientation). Every PCR check used Erm-out as the forward primer and a strain-specific reverse primer.

## Supplementary Figure Legends

**Supplementary Figure 1:**
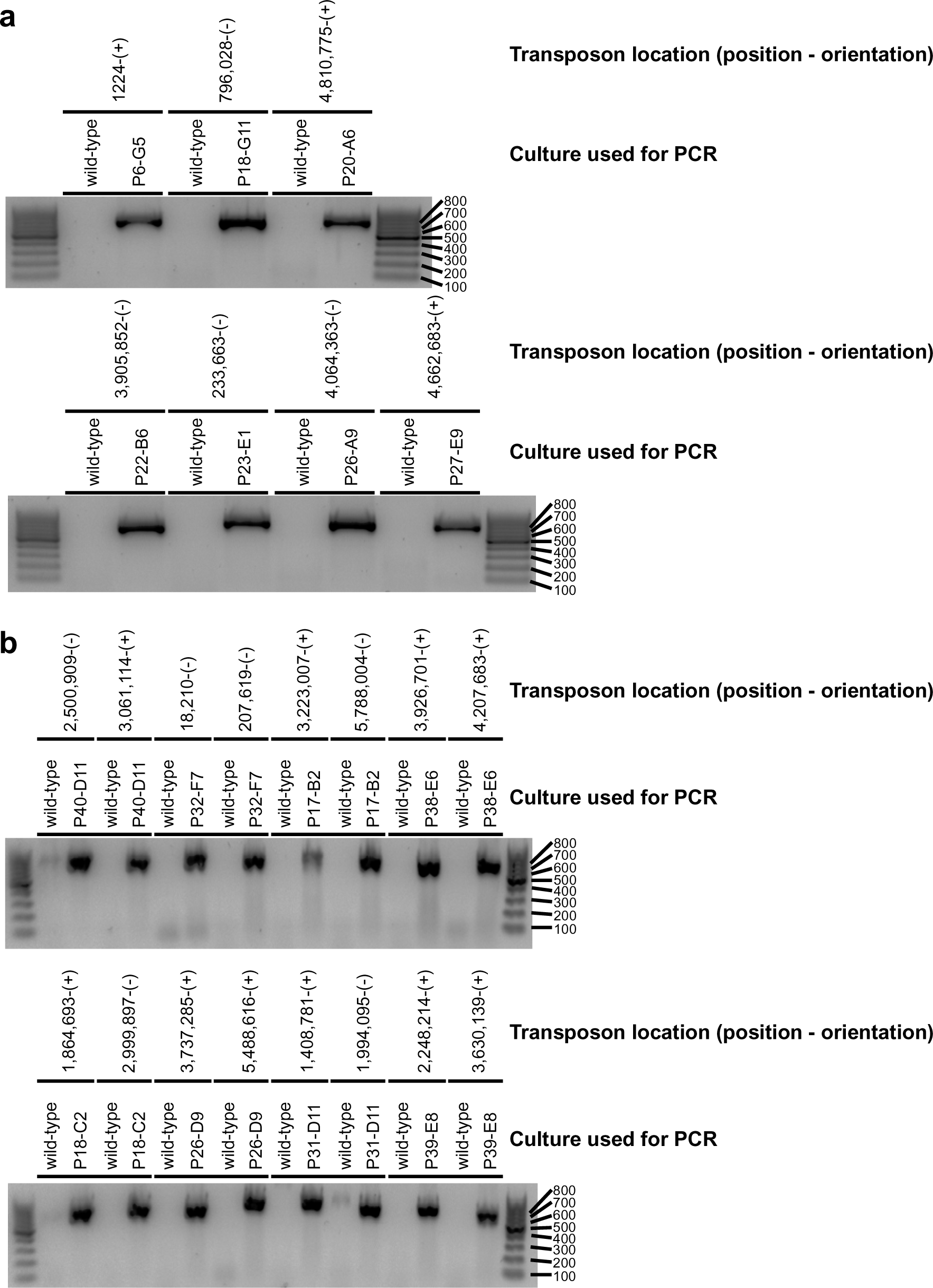
PCR validations of 15 transposon insertion strain assignments within the ordered library. PCRs were used to confirm that the protocol correctly assigns transposon insertion strains to locations in the ordered library. PCRs were performed with a universal outward facing primer that bound to the Erm cassette and a specific reverse primer that bound to the genome near the insertion point. a) Seven PCR validations of wells with a single strain with a single transposon insertion. The top label denotes the transposon insertion location that was targeted using a position-(orientation) notation. The bottom label records the culture used as input in the PCR (either an overnight culture of wild-type *B. theta* VPI-5482 or an aliquot of the cryostock in the ordered library at the position indicated, with P# abbreviating plate #). All seven wells contained the assigned transposon insertion. b) Eight PCR validations of wells with a single strain with more than one insertion. Two insertion sites were checked for each well. The lanes are labeled as in (a). All eight wells contained both of the tested transposon insertions.

